# Structures of 9-1-1 DNA checkpoint clamp loading at gaps from start to finish and ramification to biology

**DOI:** 10.1101/2023.05.03.539266

**Authors:** Fengwei Zheng, Roxana E. Georgescu, Nina Y. Yao, Michael E. O’Donnell, Huilin Li

**Affiliations:** Department of Structural Biology, Van Andel Institute, Grand Rapids, Michigan, USA; DNA Replication Laboratory, The Rockefeller University, New York, New York, USA; Howard Hughes Medical Institute, The Rockefeller University, New York, New York, USA

## Abstract

Recent structural studies show the Rad24-RFC loads the 9-1-1 checkpoint clamp onto a recessed 5′ end by binding the 5′ DNA on Rad24 at an external surface site and threading the 3′ ssDNA into the well-established internal chamber and into 9-1-1. We find here that Rad24-RFC loads 9-1-1 onto DNA gaps in preference to a recessed 5′ DNA end, thus presumably leaving 9-1-1 on a 3′ ss/ds DNA after Rad24-RFC ejects from the 5′ gap end and may explain reports of 9-1-1 directly functioning in DNA repair with various TLS polymerases, in addition to signaling the ATR kinase. To gain a deeper understanding of 9-1-1 loading at gaps we report high-resolution structures of Rad24-RFC during loading of 9-1-1 onto 10-nt and 5-nt gapped DNAs. At a 10-nt gap we captured five Rad24-RFC–9-1-1 loading intermediates in which the 9-1-1 DNA entry gate varies from fully open to fully closed around DNA using ATPγS, supporting the emerging view that ATP hydrolysis is not needed for clamp opening/closing, but instead for dissociation of the loader from the clamp encircling DNA. The structure of Rad24-RFC–9-1-1 at a 5-nt gap shows a 180° axially rotated 3′-dsDNA which orients the template strand to bridge the 3′- and 5′- junctions with a minimum 5-nt ssDNA. The structures reveal a unique loop on Rad24 that limits the length of dsDNA in the inner chamber, and inability to melt DNA ends unlike RFC, thereby explaining Rad24-RFC’s preference for a preexisting ssDNA gap and suggesting a direct role in gap repair in addition to its checkpoint role.

## INTRODUCTION

A human cell accrues up to 100,000 DNA lesions a day (1). Left unrepaired, these lesions cause mutations and programmed cell death at the cell level and aging and cancers at the organismal level (2-4). Fortunately, cells have evolved multiple DNA damage response mechanisms to detect and repair DNA damage and maintain genome integrity. One such mechanism is the cell-cycle checkpoint that encompasses an upstream damage signaling 9-1-1 DNA clamp, the associated clamp loader Rad24-RFC, and the downstream ATR–Chk kinases (5-10). 9-1-1 transduces damage signals to these kinases to arrest cell cycle progression from G1 to S during DNA replication in the S phase, or from G2 to M, in order to allow time for damage repair (11-14). The 9-1-1 clamp is a heterotrimer composed of Rad9– Hus1–Rad1 in human and Ddc1–Mec3–Rad17 in *Saccharomyces cerevisiae* (*S.c.*) (15). 9-1-1 has a ring-like structure with a pseudo-sixfold symmetry that resembles a replicative DNA clamp (16-19). However, all replicative DNA clamps are homo-oligomers, such as the homo-trimeric archaeal and eukaryotic PCNA (20-22), the homo-dimeric *E. coli* β clamp (23), and the homo-trimeric T4 phage gp45 clamp (24). The DNA replication sliding clamps stimulate activity and confer high processivity to their associated DNA polymerases and are also utilized by many other DNA metabolic enzymes (16, 25-27).

The canonical eukaryotic clamp loader is the RFC (<u>R</u>eplication <u>F</u>actor <u>C</u>) heteropentamer that loads PCNA clamps onto DNA (28-30). However, eukaryotes also contain three alternative clamp loaders that contain four subunits in common with RFC, but replace the Rfc1 subunit with another subunit, Egl1, Ctf18 or RAD17 (Rad24 in *S.c*.) (31). The Rad24(RAD17)-RFC is distinct from the other clamp loaders, as it functions with the 9-1-1 clamp instead of PCNA. The PCNA and 9-1-1 DNA clamps are topologically closed and need to be opened by ATP-driven clamp loaders to encircle the DNA duplex (25, 32, 33). The clamp loaders all have a similar two-tiered shallow spiral architecture, with five collar domains forming the top tier and five AAA+ modules forming the second tier that binds the C-terminal face of DNA clamps in the presence of ATP (24, 34-37). It has been established that RFC loads PCNA onto the DNA 3′-end by binding the DNA inside the ATPase chamber formed by the 5 subunits (24, 28-30, 38), while Rad24-RFC loads 9-1-1 onto the 5′-end (39-41), by binding the 5′ DNA at an external site on the “shoulder” of Rad24 (42, 43). Therefore, the two clamp loaders appear to use different DNA binding sites to load their respective clamps onto the opposite ends of DNA.

In light of Rad24-RFC binding a recessed 5′ end, three recent studies have revealed that RFC too has an external 5′-end DNA binding site on the external “shoulder” of Rfc1, comparable to the external site on Rad24 of Rad24-RFC, and that the external site is used for loading PCNA onto a nicked or a gapped DNA substrate for DNA repair (28-30). This has prompted us to question whether Rad24-RFC loads 9-1-1 onto a gapped DNA, as the previously published Rad24(RAD17)-RFC studies focused on 9-1-1 loading onto an isolated recessed 5′ end, and not a gap containing both 3′ and 5′ ends (40-44). The structure indicated that if 9-1-1 were loaded at gaps of around 10-nt it would place 9-1-1 onto a 3′ primer junction for possible use by TLS polymerases which biochemical studies indicate to occur (45, 46).

We find here that 9-1-1 is loaded onto gaps more efficiently than at an isolated recessed 5′ end, indicating that gaps formed during replication of damaged DNA could be used for 9-1-1 loading. However, Rad24-RFC is known to be unable to load 9-1-1 at a DNA nick, unlike RFC that can load PCNA at nicks and any gap size (28, 29, 41, 47). To understand the gap size limitation of Rad24-RFC, we designed DNAs having either a 5-nt or 10-nt gap and assembled in vitro the Rad24-RFC–9-1-1–DNA ternary complex onto each gapped DNA in the presence of ATPγS. Our cryo-EM study of the in vitro assembled complexes captured multiple 9-1-1 loading intermediates at an average resolution better than 3 Å and revealed key structural features in Rad24-RFC that underlie the preexisting gap requirement for efficient 9-1-1 clamp loading. Overall, the study supports that 9-1-1 is preferentially loaded at larger gaps and could therefore be predisposed to loading 9-1-1 clamp onto 3′ DNA for both direct DNA repair and for ATR activation.

## RESULTS

### 1. 9-1-1 loading is more efficient at a gap than at a single 5′ end

Recent structural studies of RFC reveal details of how it can load PCNA onto nicked DNA and various gaps (28-30). However, 9-1-1 clamp loading has focused on 5′ recessed ends having a 3′ ssDNA overhang with no gap and has not studied efficiency of 9-1-1 loading at a gap versus an isolated 5′ end (39, 41-43). Thus, we examined whether Rad24-RFC can load 9-1-1 onto gapped DNA and compared the efficiency to 9-1-1 clamp loading at an isolated 5′ end. For this assay we utilized two primer oligos that hybridize to a longer template oligo to form a DNA having either a 10-nt gap or isolated 5′ end (**Fig. 1a, Supplementary Table 1**). The template strand contained biotin at one end and DIG (digoxigenin) at the other, such that the DNA could be bound to streptavidin-magnetic beads at one end and the Fab of a DIG antibody at the other end, to block 9-1-1 sliding off DNA. To follow 9-1-1 clamp loading, we tagged 9-1-1 with a 7 residue PKA kinase motif and used PKA kinase to label it using α^32^P-ATP. The ^32^P-9-1-1 clamp loading reaction is performed using Rad24-RFC and magnetic DNA-bead conjugates, and then the magnetic beads were washed to remove ^32^P-9-1-1 not bound to DNA before quantitation by liquid scintillation. The results show that ^32^P-9-1-1 is loaded onto a 10-nt gap more efficiently than at an isolated 5′ end (**Fig. 1b**). Reactions lacking DNA, Mg^++^ or Rad24-RFC gave no significant ^32^P-9-1-1 loading, consistent with active ATP dependent 9-1-1 clamp loading onto the DNA by Rad24-RFC (**Fig. 1b**).

**Fig. 1.**
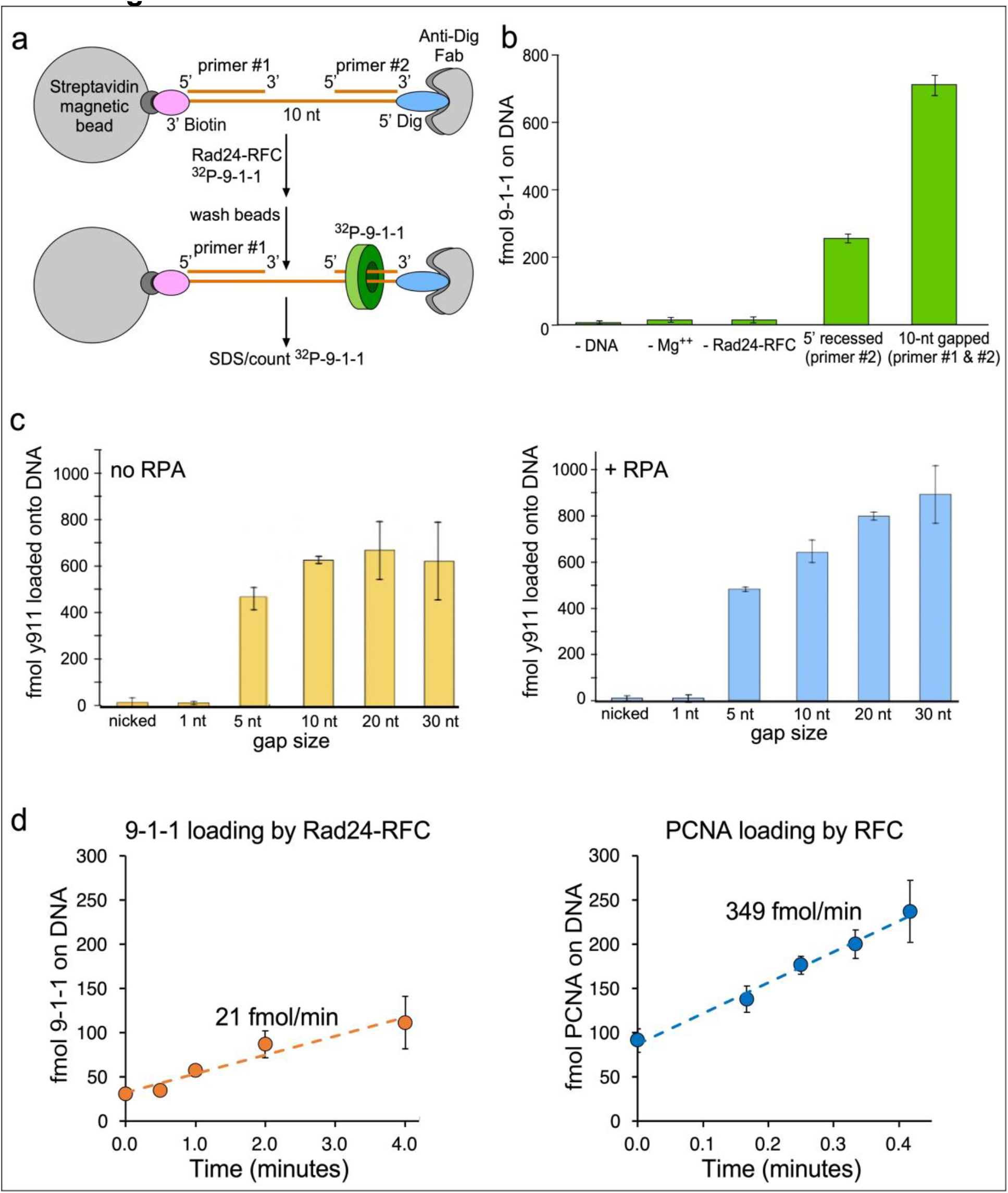
9-1-1 loading onto gapped or 5′-recessed DNA substrates. **a**) DNA oligos were annealed with a longer template DNA to provide either an isolated 5′ end (primer #2 only), or a ssDNA gap of various length (both primers #1 and #2; see Supplementary Table 1). The ^32^P-9-1-1 clamp is blocked from sliding by the biotin (attached to magnetic streptavidin beads) on one end and the DIG moiety attached to the Fab of an antibody to DIG at the other end. See **Methods** for details. **b)** Comparison of 9-1-1 loading at a recessed 5’ end and at a 10 nt gap in the presence of RPA, along with control reactions lacking one assay component. **c)** Magnetic bead assays using nicked DNA or different sized gaps either minus RPA (left) or plus RPA (right). **d**) Comparison of DNA loading rates of 9-1-1 (by Rad24-RFC) and PCNA (by RFC) were performed under identical conditions using the 10-nt gap DNA. All experiments in panels b, c and d were performed in triplicate and the error bars represent standard error of the mean.

While RFC can load PCNA at nicks and any gap size, Rad24-RFC has been shown to be incapable of loading 9-1-1 at a nick (28, 29, 41, 47). Hence, we examined ability of Rad24-RFC to load 9-1-1 onto gaps of different sizes. The results show that while Rad24-RFC does not load 9-1-1 at a nick or a 1-nt gap, it is capable of loading 9-1-1 onto gaps of 5-30 nt (**Fig. 1c**). The presence of RPA did not change the results significantly but appeared to stimulate the reaction somewhat as the gap size increased. Considering that a 1-nt gap is not utilized by Rad24-RFC/9-1-1, one may conclude that short base excision gap repair does not utilize 9-1-1. However, 9-1-1 loading at gaps of 5 nt or more indicates that 9-1-1 may be utilized in long base excision gap repair, or nucleotide excision repair (see Discussion).

When the 9-1-1 clamp is loaded onto 5′ ends it signals the cell cycle checkpoint, but numerous 5′ ends are formed during normal lagging strand replication in the absence of damage, occurring every 100-200 bp (48). Thus, one may question how the cell prevents Rad24-RFC from utilizing these numerous 5′ sites that could potentially result in cell cycle arrest? To address this, we compared the kinetics of RFC loading of PCNA onto DNA and Rad24-RFC loading of 9-1-1 onto DNA. The results show that RFC/PCNA loading is an order of magnitude more rapid than Rad24-RFC–9-1-1 (**Fig. 1d**). Therefore, we propose that during normal replication Okazaki fragments are filled-in and ligated faster than 9-1-1 loading could occur. This proposal is based in recent elegant studies (49) that indicate Pol δ/PCNA/Fen1/ligase is exceedingly rapid in completing Okazaki fragment maturation. Therefore Rad24-RFC may not have time to load 9-1-1 onto 5′ ends of lagging strand fragments during unobstructed DNA replication.

### 2. Cryo-EM structure of Rad24-RFC–9-1-1 assembled on a 10-nt gapped DNA substrate

To examine the structure of 9-1-1 loading onto a 10-nt gapped DNA, we mixed separately purified Rad24-RFC and 9-1-1 clamp with a 10-nt gapped DNA substrate (**Fig. 2a**) at a molar ratio of 1.0:1.2:1.6 and in the presence of 0.5 mM weakly-hydrolysable ATP analog ATPγS and 5 mM Mg^2+^ (see **Methods** for details). The mixture was used directly to make cryo-EM grids without further purification. 2D class averages of selected particle images showed successful reconstitution of the Rad24-RFC–9-1-1 clamp– 10-nt gapped DNA complex (**Supplementary Fig. 1a**). After multiple rounds of 2D and 3D classifications, 3D reconstruction, and non-uniform refinement by using cryoSPARC (50) and Relion (51), we obtained five EM maps. These maps had a similar dimension of 129 Å × 113 Å × 108 Å and had an average resolution range of 2.76 Å to 2.94 Å (**Supplementary Figs. 2-3**).

**Fig. 2.**
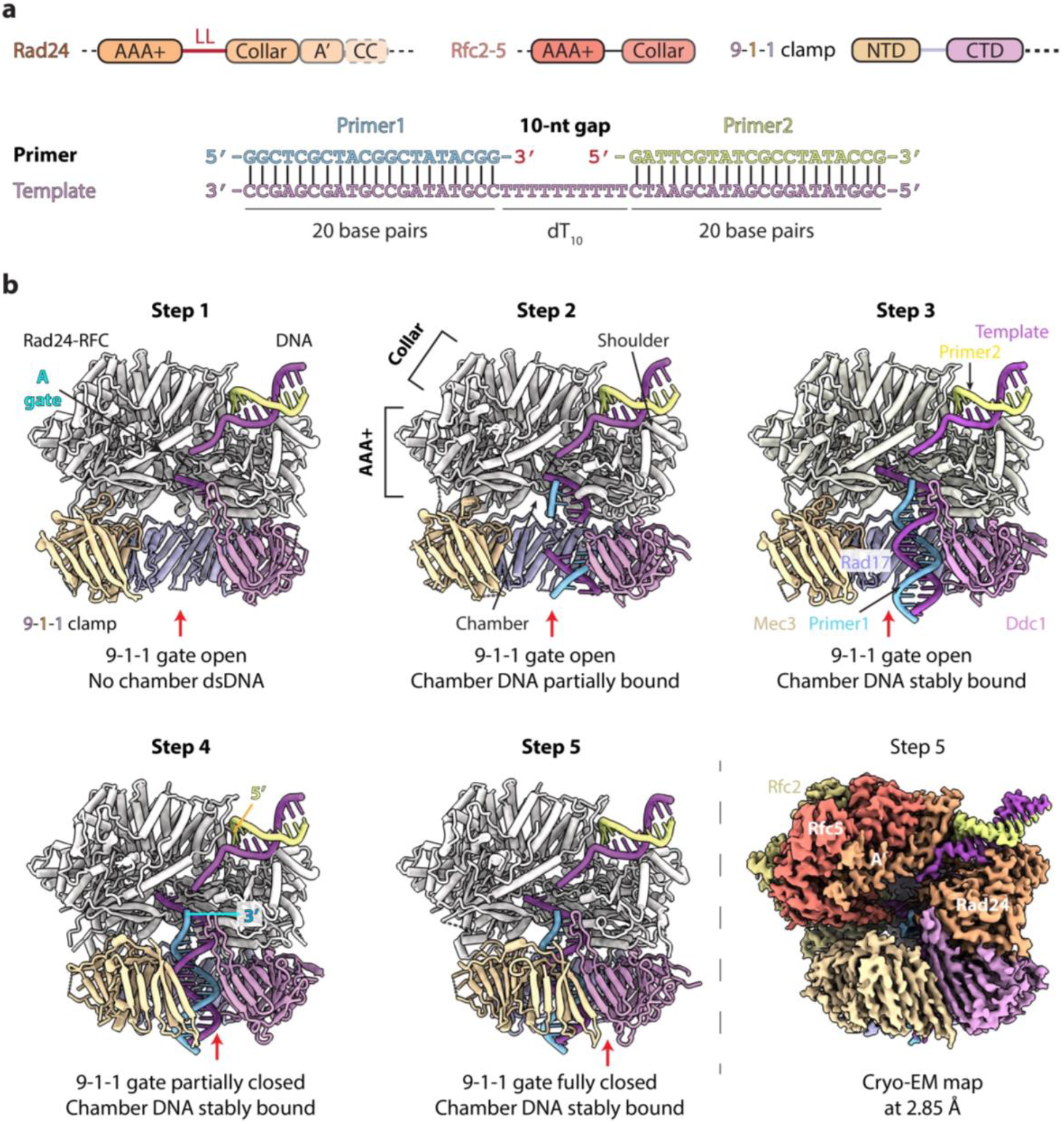
Five Rad24-RFC intermediates in loading 9-1-1 clamp onto a 10-nt gapped DNA. **a**) Domain architecture of Rad24-RFC, 9-1-1 clamp, and the 10-nt gapped DNA. The Rfc2-5 and three subunits of 9-1-1 have similar respective domain arrangement and are shown together for brevity. Dashed lines indicate unsolved regions in EM maps. LL is the long linker between Rad24 AAA+ module and collar domain. NTD and CTD in each 9-1-1 subunit are linked by the inter-domain connecting loop (IDCL). The 10-nt gapped DNA harbors both 5′ and 3′ junctions. **b**) Structures of five Rad24-RFC loading intermediates of 9-1-1 clamp, arranged in a plausibly temporary sequence based on the progression of DNA binding in the central chamber of Rad24-RFC. In step 1, the 9-1-1 gate is open, and the DNA has not bound into the clamp loader chamber. In step 2, the 9-1-1 gate remains open, the DNA has entered the central chamber and passed through the 9-1-1-gate, but the DNA is partially stable, and the EM density for 4-bp DNA between the loader and the clamp are missing. In step 3, the 9-1-1 remains open, and the DNA in the chamber is fully stabilized. In step 4, the 9-1-1 gate is partially closed, and DNA is fully engaged. In step 5, the 9-1-1 gate is fully closed around the loaded DNA. A representative cryo-EM density map rendered at a high threshold (0.2) is shown at the lower right corner. For clarity, the Rad24-RFC structure in all five models is in ivory. The red arrow points to the 9-1-1 gate. The DNA entry A-gate in the Rad24-RFC is labeled in the step 1 structure. The structures are aligned and shown in the same front view.

We built atomic models for all five maps based on published structures of Rad24-RFC–9-1-1 bound to a 5′ tailed DNA (42, 52) and refined these models to < 3 Å resolution (**Supplementary Table 2**). The overall architecture of the five Rad24-RFC–9-1-1–10-nt gapped DNA complex structures are similar to previously reported clamp–clamp loader complexes (24, 35, 36, 38, 42, 52). They all adopt the expected three-tiered architecture, with Rad24-RFC forming the top (collar domains) and middle (AAA+ modules) tiers that sit above 9-1-1 at the bottom tier (**Fig. 2a-b**). The 10-nt gapped DNA used in this study contains three regions: the 3′ ss/ds junction (primer1/template DNA), the middle 10-nt ssDNA gap (residues 21-30 nt of the template strand), and the 5′ ss/ds junction (primer2/template DNA) (**Fig. 2a**). In all five structures, the 5′ ss/ds junction DNA is stably bound to the external shoulder of Rad24, and the Rad24-RFC loader is highly similar, with a root-mean-square deviation (RMSD) between main chain C_α_ atoms ranging from 0.6 Å to 0.7 Å (**Video 1**).

Logically, these structures represent the Rad24-RFC–9-1-1 clamp–10-nt gapped DNA complex in five different DNA binding and clamp loading stages: we assigned the 2.93 Å resolution structure with an open 9-1-1 clamp but without the 3′ ss/ds junction DNA as the step 1 intermediate; the 2.94 Å resolution structure with an open 9-1-1 and partially bound 3′ ss/ds junction DNA as the step 2 intermediate; the 2.76 Å resolution structure with an open 9-1-1 and with stably bound 3′ ss/ds junction DNA density as the step 3 intermediate; the 2.90 Å resolution structure with a gate partially closed 9-1-1 as the step 4 intermediate; and the 2.85 Å resolution structure with a gate fully closed 9-1-1 encircling the 3′ ss/ds junction DNA as the step 5 intermediate (**Fig. 2b**).

### 3. The Rad24 upper loop blocks the 3′-ss/ds junction and sets the minimum ssDNA gap size

Rad24-RFC recognizes the DNA within its internal chamber (i.e. 3′ ss/ds junction DNA) in a similar manner as the yeast RFC (29, 30, 38). The α-helices 4 and 5 (α4 and α5) in the AAA+ modules of all Rad24-RFC subunits wrap around and trace the template strand of 3′ ss/ds junction DNA, forming H-bonds with the template phosphate backbone (**Fig. 3b**). For example, Rad24 Met-163, Rfc5 Arg-106, Rfc4 Arg-90, Rfc3 Ile-90, and Arg-94, and Rfc2 Ile-103 and Arg-107 of α4-helices form H-bonds with the template strand, and Rfc3 Thr-123 of α5 helix forms an H-bond with the template strand. The loader has only very limited interactions with the 3′ primer strand (i.e., Rad24 Arg-199 of α5 and Asn-80 of the Rfc5 plug form H-bonds with that strand), see also **Supplementary Fig. 4c**.

**Fig. 3.**
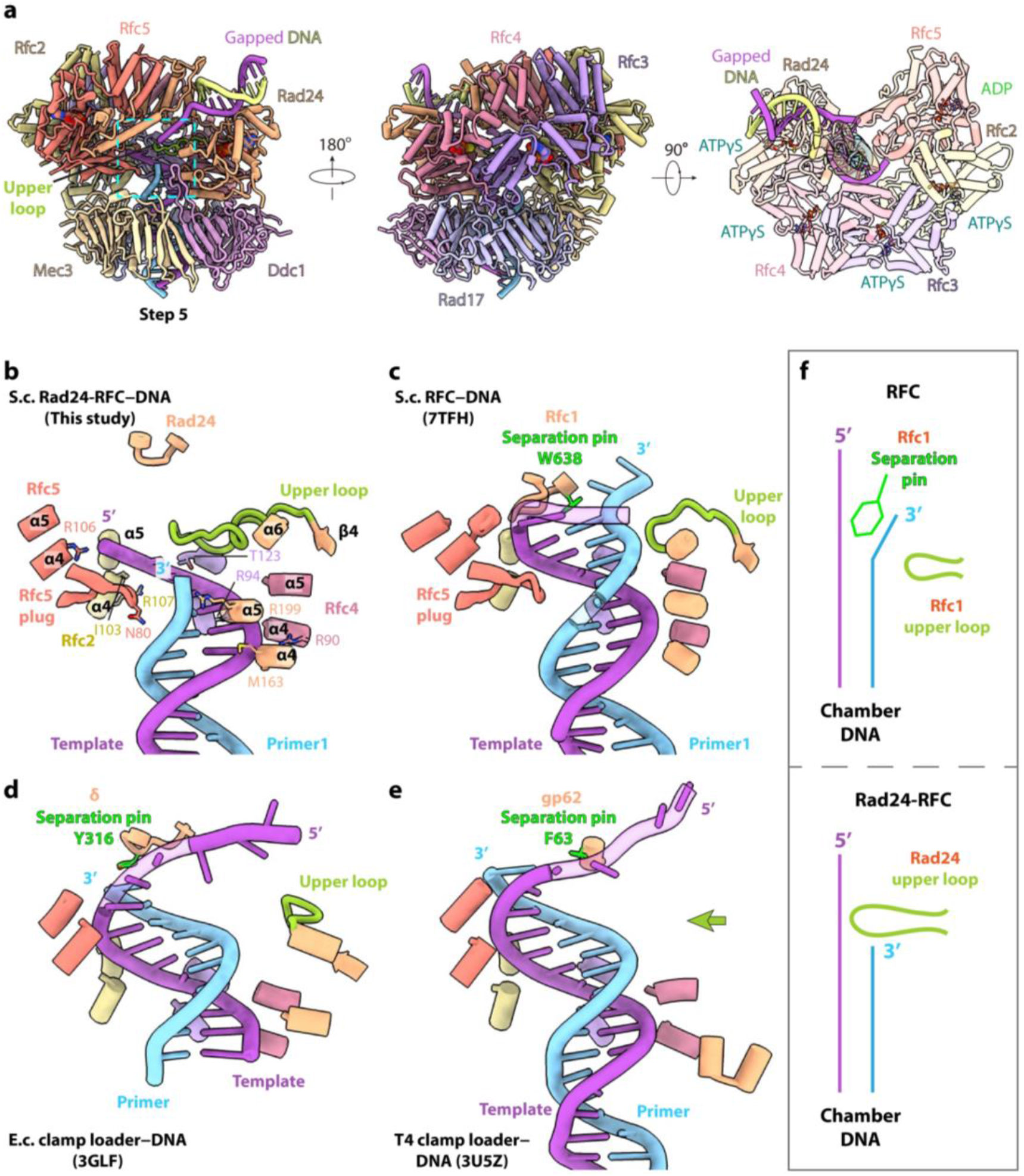
The Rad24 upper loop sets the gap size between the 3′ and 5′ DNA junctions. **a**) The step-5 structure of Rad24-RFC–9-1-1–10-nt gapped DNA in a front (left), back (middle), and top (right) view. Subunits are individually colored. 9-1-1 is omitted in the right panel to better show the bound ATPγS and ADP. The Rad24 upper loop inside the chamber is highlighted in green. The region in the cyan box is shown enlarged in b. **b-e**) Comparison of DNA binding in the chamber of Rad24-RFC (b), RFC (c), *E. coli* clamp loader (d), and T4 phage clamp loader (e). **f**) Sketch comparing the 3′ DNA binding mode in the chamber of RFC and Rad24-RFC based on structures in panels **b** and **c**. The loaders are aligned but omitted except for a few labeled key elements. The α4 and α5 helices of each loader subunit follow and wrap around the template strand in purple. In (b), Rad24 Arg-199 and Rfc5 Asn-80 H-bond with the primer phosphate backbone in blue. The Rad24 upper loop blocks the primer strand from advancing upward, in contrast to the higher reach of the primers in all other loaders. The equivalent loops in other loaders are much shorter and do not block the primer strand (**c-d**). The T4 loader lacks the equivalent loop (indicated by a green arrow) as the corresponding AAA+ module is highly degenerated **(e)**, also see **Supplementary Figs. 4b and 5**. Notably, all clamp loaders − except for Rad24-RFC − harbor an aromatic residue (lime) at the top that functions as a separation pin to unwind DNA from the 3′-junction.

In the previously reported DNA bound structures (42, 43), there were two structural elements in Rad24-RFC that appeared to be narrowing the DNA path: a β-hairpin coming from Rcf5 (also termed RFC5 E-plug), and a long loop positioned at the upper part of the inner chamber coming from Rad24 (termed upper loop). We therefore hypothesized that these structural elements might be responsible for the unique DNA binding behavior of Rad24-RFC. In this study with the 10-nt gapped DNA, we found that a dsDNA segment – the 3′ ss/ds DNA – can actually be accommodated into the lower chamber of Rad24-RFC, with the 3′-junction residing just below the upper loop (**Fig. 3a**).

Interestingly, we found that it is the Rad24 upper loop that is primarily responsible for blocking the 3′-junction, thereby preventing the duplex region of the DNA from advancing further up the central channel (**Fig. 3b, Video 2, Supplementary Fig. 4a**). The Rfc5 E-plug can accommodate dsDNA and apparently plays an accessory role to the Rad24 long loop. Of the 10-nt poly(dT) ssDNA gap used in our DNA substrate, up to seven nucleotides could be modeled based on electron density: in both steps 3 and 5 structures, we modeled four nucleotides adjacent to the 3′-ss/ds DNA junction and three nucleotides near the 5′-ss/ds DNA junction, the 22-Å density gap can be connected by three additional nucleotides, accounting for the total 10-nt poly(dT) ssDNA in the template strand DNA (**Fig. 4a** bottom middle). A 9-nt ssDNA distance between a sensor chip and a 5′ end was previously shown to enable 9-1-1 loading by Rad24-RFC, consistent with the current study (41).

**Fig. 4.**
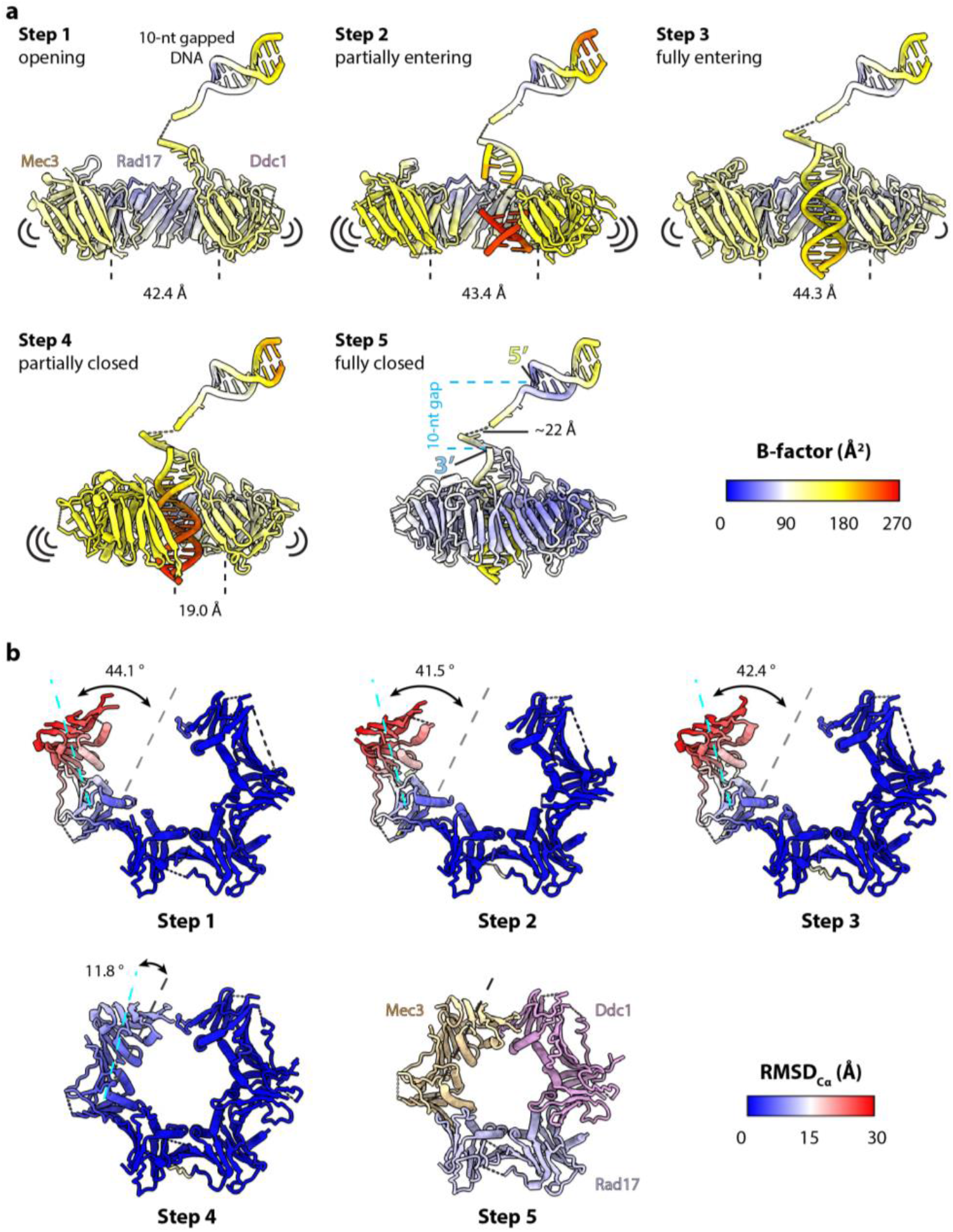
Gradual closure of the 9-1-1 gate accompanying DNA binding. **a**) Flexibility analysis of the five loading intermediates. The atomic models are colored by their respective local b-factors in ChimeraX. Rad24-RFC is omitted for clarity. Mec3 and Ddc1 lining the DNA gate are partially mobile (yellow) in the open gate and partially open gate in steps 1-4, but they become stable (blue) in the closed gate in step 5. Consistent with the assigned temporal sequence, the chamber DNA is absent in step 1, is present but flexibly bound (red) in step 2 and becomes more stably bound (yellow) in step 3. The partial gate closure in step 4 destabilizes the chamber DNA (red), likely due to the perturbation by Mec3 movement. The chamber DNA is better stabilized in the 9-1-1 gate fully closed step 5. The 9-1-1 gate size is labeled. Four nucleotides and three nucleotides near the top 5′-junction and the bottom 3′-junction, respectively, are stabilized in the Rad24-RFC chamber. The dashed line represents three disordered nucleotides in the gap region. Because the Rad24-RFC A-gate is open in all structures, ssDNA longer than 10-nt in the gap region can be easily accommodated by looping outward through the A-gate. **b**) Although both Mec3 and Ddc1 line the DNA entry gate, Mec3 is the actual “gate” of the 9-1-1 clamp. The 9-1-1 structures in steps 1-4 are colored by their local RMSD_Cα_ compared to the step 5 structure. Ddc1 is static, and gate opening and closing only involve in-plane rotation of Mec3.

### 4. Explanation for inability of Rad24-RFC to unwind DNA like the canonical RFC

RFC was recently shown to harbor a separation pin (Rfc1 Trp-638) to unwind dsDNA from a 3′-ss/ds DNA junction (29, 38), and is capable of unwinding a nicked dsDNA to form a 6-nt ssDNA gap for PCNA loading using ATPγS, achieved by clamp loader-to-DNA binding energy and not requiring hydrolysis (28). A side-by-side comparison of the Rad24-RFC–9-1-1–DNA structure (step 5) with several structures of the replicative clamp loader–clamp–DNA complexes reveals two striking distinctions (**Fig. 3b-f, Supplementary Figs. 4b and 5**). First, the upper loop of Rad24 is much longer than corresponding loops of the *S. cerevisiae* RFC and *E. coli* clamp loader, and the T4 clamp loader lacks an equivalent loop. Second, while all replicative clamp loaders possess a DNA duplex separation pin (i.e. Trp-638 in yeast Rfc1, Tyr-316 in *E. coli* δγ_3_δ′, and Phe-63 in T4 phage gp44/62 clamp loader) (24, 29, 53), a similar duplex separation pin is absent in Rad24. Further, the Rad24 loop that lacks the separation pin is located much higher within the central chamber than the corresponding loops in the replicative clamp loaders harboring the separation pin. By superimposition, we found that the 3′-junction of primer/template in all replicative loader complexes (RFC, *E. coli* δγ_3_δ′, and T4 loader) would clash with the Rad24 long upper loop (**Fig. 3c-e, Supplementary Figs. 4b and 5**). Based on these observations, we suggest that Rad24-RFC has evolved the long upper loop to specifically stop the 3′-ss/ds primer template junction at just below this loop, thereby enforcing a minimum ssDNA gap size (i.e., ssDNA region in the template strand between the lower 3′-junction and the upper 5′-junction of the gapped DNA). We further suggest that because the 3′-junction is blocked in Rad24-RFC and no longer requires unwinding, Rad24 has lost the separation pin during evolution. Consistent with the presence of 3′-junction unwinding activity of the replicative clamp loaders and the absence of an unwinding activity of Rad24-RFC (29, 38-41), there is a side tunnel in yeast RFC from which the unwound primer 3′-tail emerges, but this tunnel is narrower and is blocked at the bottom by the Rad24 upper loop in Rad24-RFC (**Supplementary Fig. 6a-b**).

### 5. Five intermediate structures reveal a top-to-bottom binding process in loading 9-1-1 onto a 10-nt gapped DNA

As mentioned above, in those five 10-nt gapped DNA bound intermediates, the upper loop of Rad24-RFC interacts with 5′ ss/ds junction DNA similarly; the main differences lie in the size of the DNA entry gate of 9-1-1 and the binding of the 3′ ss/ds junction DNA in the central chamber of the loader (**Fig. 4a**). In the step 1 structure, the 9-1-1 DNA gate is 42 Å wide, and only four nucleotides in the 10-nt ssDNA gap region near the 3′-junction are visible. In the step 2 structure, the 9-1-1 gate remains wide open. There is relatively weak DNA density in the loader chamber as well as inside the 9-1-1 clamp, and the density is broken at the interface between the loader and the clamp. In the step 3 structure, the chamber DNA is stably bound with strong density (P1 6-20 and T 31-45 with a 15-bp dsDNA region), although the 9-1-1 gate is still open. In the step 4 structure, the 9-1-1 gate starts to close around the stably bound 3′ ss/ds junction DNA in the chamber of the loader as the gate has narrowed to 19 Å. In contrast, the 9-1-1 gate is totally closed around the DNA in the step 5 structure (**Fig. 4a**).

It is clear from the above description that the DNA binding is a three-stage top-to-bottom process that starts with the 5′-ss/ds junction binding to the external shoulder site of Rad24-RFC on the top. This is followed by the gap ssDNA region and the top region of the 3′-ss/ds DNA junction DNA entering the inner chamber of Rad24-RFC through the A-gate (**Fig. 2b,** upper left) that is open between the Rad24 N-terminal AAA+ module and C-terminal A′ domain (step 1). And finally, the 3′ ss/ds DNA region enters the 9-1-1 central channel (steps 2-3).

### 6. 9-1-1 gate closure accompanies the binding of the 3′ chamber DNA

We next performed a flexibility analysis of the five loading intermediates, to understand the relationship between DNA binding and 9-1-1 gate closure. We used the B-factors of the atomic models, a measurement of thermal motion, as an indicator of the structural flexibility (**Fig. 4a**). In the step 1 structure, the 5’ ss/ds junction DNA at the external shoulder site of Rad24 is stably bound except at the terminus (orange), and the gate-lining subunits Ddc1 and Mec3 of 9-1-1 are partially flexible (light yellow), but the third 9-1-1 subunit (Rad17) is stable (light blue). The 42-Å wide gap in 9-1-1 is more than sufficient to allow passage of the 20 Å wide duplex of the 3′ ss/ds junction DNA into the central chamber of the loader. In the step 2 structure, both 3′ ss/ds junction DNA in the chamber (orange to red) and 9-1-1 (yellow) become more flexible as the 3′ ss/ds junction DNA is entering the clamp. This is compared to the step 3 structure in which both DNA and the 9-1-1 gate become stabilized upon completion of DNA entrance in step 2. Interestingly, in the step 4 structure, the DNA gate is only 19 Å wide, and not only the gate lining Ddc1 and Mec3 (yellow) but also the 3′ ss/ds junction DNA (red) in the chamber becomes flexible. This is probably due to the perturbation by Mec3 movement to close the 9-1-1 gate. As expected, both 9-1-1 (light blue) and DNA (light yellow to light blue) are stabilized in the DNA fully bound and gate fully closed step 5 structure. Therefore, we conclude that 9-1-1 gate closure progresses along with the gradual insertion and eventual stable binding of the 3′-ss/ds junction DNA in the chamber. Taken together, we suggest that the five intermediates captured by cryo-EM can be considered as a temporal and sequential loading process of the 9-1-1 clamp onto a gapped DNA by Rad24-RFC.

### 7. Mec3 functions as the DNA entry gate of the 9-1-1 clamp

We noticed that both gate-lining subunits Mec3 and Ddc1 are flexible during the gate closure process, but the flexibility of Mec3 is significantly more pronounced than Ddc1 (**Fig. 4a**), particularly in the step 4 structure where the gate has partially closed (compare the yellow-to-orange Mec3 with the light yellow Ddc1), and in the step 5 structure with a fully closed 9-1-1 gate (compare the white Mec3 with the blue Ddc1). This observation implies that gate closure may be asymmetric in that the two gate subunits contribute unequally to the clamp closing process. To investigate this more quantitatively, we superimposed the 9-1-1 clamps in steps 1-4 with the gate-closed 9-1-1 clamp in step 5, and calculated their respective RMSD_Cα_, and colored the clamps by their respective RMSD_Cα_ values (**Fig. 4b**). The results show that the 9-1-1 gate opening is mediated by an in-plane ∼42° rigid-body rotation of Mec3 pivoting on the interface region between Mec3 and Rad17 (steps 1-3). Therefore, the partial gate closure in step 4 is accomplished by Mec3 rotating back towards Ddc1 by 30°. The final gate closure from step 4 to step 5 involves another 12° rotation of Mec3 to contact Ddc1 (**Fig. 4b**). Therefore, we suggest that Ddc1 is largely stationary, and the 9-1-1 gate opening and closure is primarily mediated by a rigid-body rotation of Mec3.

### 8. The 9-1-1 clamp ring remains flat during gate opening and closure

The DNA clamp studies thus far are only flat rings when alone or encircling dsDNA (31). However, during loading of DNA replication clamps onto DNA, the clamp is cracked open by the clamp loaders and forms a lock washer like spiral structure, exemplified by the PCNA lock washer being loaded onto DNA by RFC (28, 29, 47), as well as the T4 clamp lock washer being loaded onto DNA by the T4 clamp loader (24, 33) (**Supplementary Fig. 7**). However, in all five steps of 9-1-1 clamp loading onto DNA by Rad24-RFC, the 9-1-1 clamp remains flat, despite its various degrees of gate opening (**Figs. 2** and **4, Video 3, Supplementary Fig. 7).** Therefore, the in-plane mechanism of gate opening/closure of the 9-1-1 clamp appears to be somewhat unique in this aspect. In fact, the 9-1-1 clamp is unique among the DNA clamps in eukaryotes in that it is the only eukaryotic ring composed of three different subunits; the bacterial and eukaryotic cellular replicative DNA clamps are either homo-dimeric or homo-trimeric and open out-of-plane. Further study is needed to understand if the heteromeric structural feature is responsible for the unique gate opening/closure mechanism of the 9-1-1 clamp.

### 9. Rad24-RFC binds DNA differently at a 5-nt gap

As explained above, the structure of Rad24-RFC suggests that the upper loop will limit the size of gap that 9-1-1 can be loaded on. Therefore, we next imaged the mixture of Rad24-RFC, 9-1-1, and a 5-nt gapped DNA (**Fig. 5a**) in the presence of ATPγS, prepared in the same condition as with the 10-nt gapped DNA. 2D and 3D classification of cryo-EM images showed that most Rad24-RFC–9-1-1 complex particles did not contain DNA (**Supplementary Fig. 1b**). For the particles with some DNA density in the central chamber of Rad24-RFC, focused refinement around DNA showed that the DNA density was either partial or distorted from the B-form structure in 4 out of the 5 subclasses. Further refinement of the subclass with undistorted and relatively strong DNA density resulted in the final 3D map at 3.04 Å overall resolution (**Supplementary Figs. 3f-g and 8**). We observed a lower percentage of DNA-loaded particles using 5-nt gapped DNA vs 10-nt gapped DNA, consistent with the somewhat lower, but repeatable extent of loading 9-1-1 onto 5-nt gap DNA compared to loading of 9-1-1 at larger gaps in Fig. 1. This observation indicates Rad24-RFC has difficulty loading 9-1-1 onto a 5-nt gapped DNA vs 10mer gapped DNA.

**Fig. 5.**
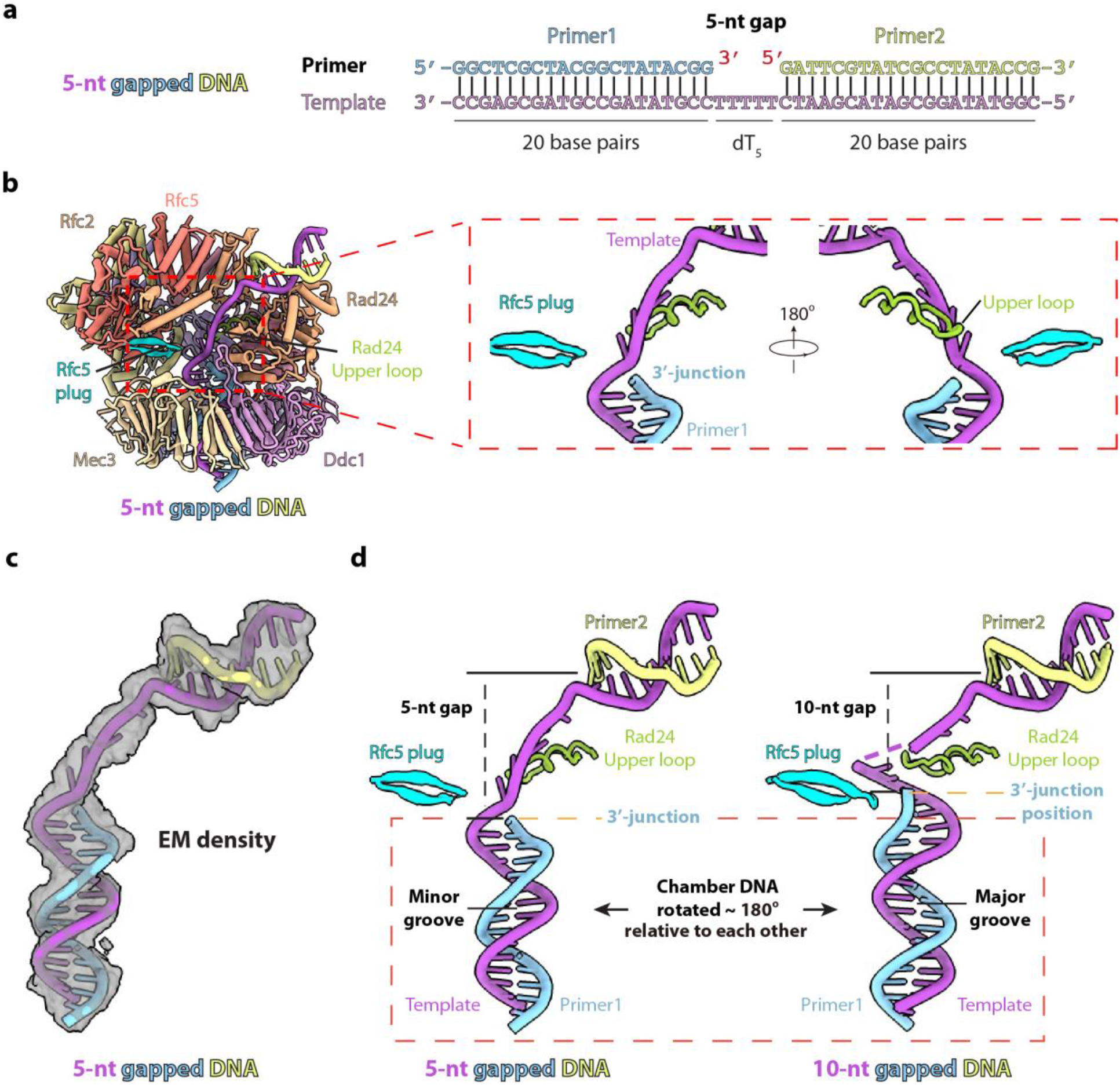
The Rad24 upper loop remains in place to block the advance of 3′ junction of the 5-nt gapped DNA. **a**) A sketch of the used 5-nt gapped DNA substrate. **b**) Structure of the Rad24-RFC–9-1-1–5-nt gapped DNA complex in a front view. Subunits are individually colored. Right panel shows enlarged views of the 5-nt DNA gap region. The Rad24 upper loop and the Rfc5 plug are shown at the 3′ DNA junction to block the upward movement of the chamber DNA (i.e., DNA bound in the central chamber of Rad24-RFC). **c**) EM map of the 5-nt gapped DNA is shown in transparent grey surface and superimposed on the atomic model. The auto-refined map by Relion before post-processing at 3.66 Å resolution has stronger DNA density and is used here. **d**) Side-by-side comparison of the Rad24 upper loop (green) and Rfc5 plug (cyan) interacting with the 5-nt gapped DNA (left) and the 10-nt gapped DNA from step 5 (right). Note that the lower 3′ dsDNA in the left structure is rotated by 180° around its helical axis, such that the purple template strand is oriented to connect with the upper 5′ dsDNA with a minimum length (5 nt).

The 5-nt gapped DNA bound Rad24-RFC–9-1-1 structure resembles the step-5 structure of the 10-nt gapped DNA bound complex. The overall structures are nearly superimposable with an RMSD of main chain C_α_ atoms of 0.6 Å, and both 9-1-1 clamps are fully closed (**Fig. 5b, Supplementary Fig. 9, Video 1**). The three intermediate steps 2-4 seen using the 10-nt gapped DNA were not observed using the 5-nt gapped DNA. Hence, it is possible that the intermediate states of the 9-1-1 ring using the 10-nt gapped DNA are nearly isoenergetic forms, stochastic in nature, and that these states are not stable at a strained 5-nt gapped DNA, resulting in only the closed form.

An important difference is observed in the 3′ ss/ds DNA within the central chamber (**Fig. 5c**). This DNA region was rotated 180° around the helical axis compared to the 10-nt gapped DNA (**Fig. 5d**). In the rotated pose, the template strand is oriented the closest to the upper 5′ shoulder DNA, enabling the short 5-nt ssDNA to bridge the gap between the two dsDNA segments. Interestingly, the Rfc5 hairpin plug now contacts the minor groove of the rotated 3’ ss/ds DNA of the 5-nt gapped DNA, distorting the DNA and in contrast with the Rfc5 plug contacting the major groove of the chamber DNA of the 10-nt gapped DNA. However, the Rad24 upper loop is in a similar position in the two structures, blocking the upward advance of the 3′ ss/ds DNA segment. Therefore, no unwinding of either the 3′ or 5′ dsDNA ends of the 5-nt gapped DNA was observed, underscoring the absence of separation pins in the Rad24-RFC clamp loader as described above. In summary, the 5-nt gapped DNA bound structure reveals a possible cooperation between the Rfc5 plug and Rad24 upper loop in controlling the minimal DNA gap size accessible to Rad24-RFC and confirms the absence of DNA unwinding activity in Rad24-RFC.

## DISCUSSION

### 1. 9-1-1 loading is more efficient on a gapped DNA vs 5′-recessed DNA

The current study shows that Rad24-RFC is more efficient in loading the 9-1-1 clamp onto a gapped DNA compared to an isolated recessed 5′ DNA terminus. Indeed, our cryo-EM structural results show that the 3′ duplex of gapped DNA binds in the central channel of Rad24-RFC, compared to exclusively ssDNA in the case of an isolated recessed 5′ DNA. It therefore seems likely that the enhanced 9-1-1 loading at gaps vs single 5′ ends may be due to Rad24-RFC binding the 5′ dsDNA at the same time as the 3′ ss/ds DNA at gaps. This geometry predicts the 9-1-1 clamp will be loaded around a 3′ primer terminus instead of ssDNA. In this case, the 9-1-1 clamp is positioned at a 3′ ss/ds DNA (similar to PCNA) and could be used by TLS Pols as indicated in earlier studies (54-56) and as described further below. We presume that most DNA lesions on the lagging strand template during replication will result in stopping Pol δ–PCNA and thus require TLS Pols or recombinative repair to bypass the lesion, extending the time of Okazaki fragment gap fill-in. Thus, lagging strand gaps might be a major source of 5′ ends for 9-1-1 loading and cell cycle arrest during DNA damage in S phase.

### 2. A proposed model for 9-1-1 loading onto a gapped DNA

Based on previous biochemical and structural studies of this system (39-42, 52), recent structural knowledge of PCNA loading by RFC (28-30, 38), and our current 9-1-1 clamp loading intermediates (steps 2-4) described above, we propose here a significantly more detailed 9-1-1 loading process, specifically onto gapped DNAs by Rad24-RFC (**Fig. 6a**). Before encountering a DNA, we suggest that the 9-1-1 ring is opened by Rad24-RFC in the presence of ATP, in a mechanism resembling the PCNA gate opening by RFC in the absence of DNA (**Supplementary Fig. 7**) (38). Consistent with this suggestion, it has been shown that in the presence of ATP or ATPγS, Rad24-RFC and the 9-1-1 clamp can form a complex without a DNA substrate (39, 40), although an experimental structure of Rad24-RFC with an open 9-1-1 ring in the absence of DNA has yet to be captured. In step 1, we suggest that Rad24-RFC first engages the 5′ duplex at the external (shoulder) DNA binding site of Rad24. This is followed by steps 2 and 3 in which the ssDNA at the gap as well as the 3′ ss/dsDNA region gains access to the Rad24-RFC central chamber and the 9-1-1 central channel to become stably bound there. We further suggest that during this top-to-bottom DNA binding process, the Rad24 upper loop plays a crucial role in selecting against binding a DNA substrate with a ssDNA gap less than 5-nt. In step 4, Mec3 rotates towards Ddc1 to close the DNA entry gate in 9-1-1. In step 5, we suggest that the 9-1-1 gate closing on the stably bound DNA will activate the ATPase activity in Rad24-RFC, leading to ATP hydrolysis, release of inorganic phosphate, and the dissociation of ADP-bound Rad24-RFC. These actions leave the closed 9-1-1 clamp encircling 3′ duplex DNA (step 6).

**Fig. 6.**
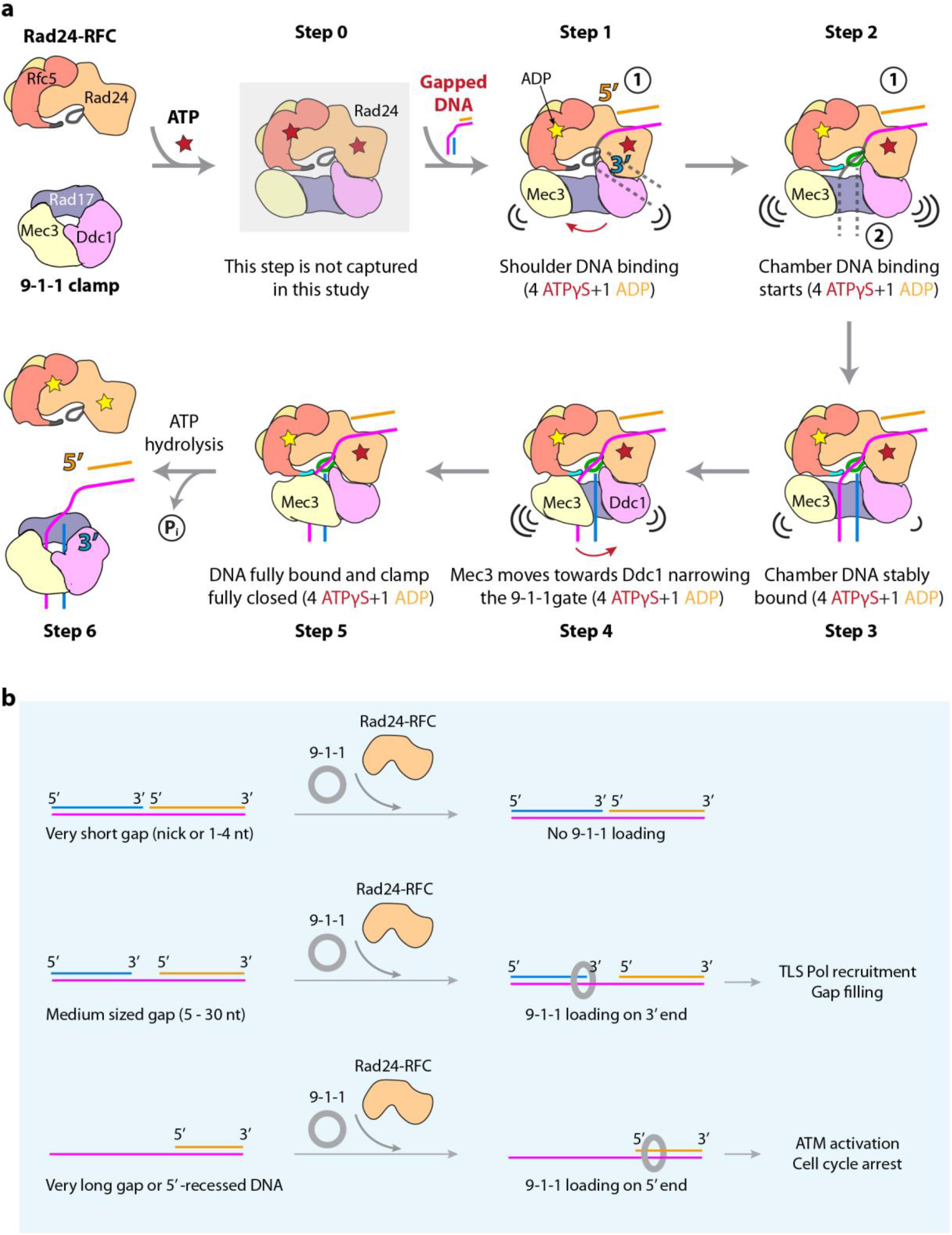
Proposed model of Rad24-RFC loading the 9-1-1 clamp onto a gapped DNA. **a**) In preparation for loading (step 0), Rad24-RFC binds the 9-1-1 clamp to form a binary complex in the presence of ATP (or ATPγS used in this study). In a mechanism similar to the mutual activation of RFC and PCNA (38), we suggest that binding energy between Rad24-RFC and 9-1-1 drives the DNA gate opening in Rad24-RFC (the A-gate) and 9-1-1 (between Mec3 and Ddc1). This intermediate is greyed out as it is yet to be captured. In step 1, the shoulder DNA binds first to the Rad24-RFC external site. This is followed by the chamber DNA binding and passing through the 9-1-1 gate in steps 2 and 3. Once the DNA has fully entered, in steps 4 and 5, Mec3 moves towards Ddc1 to close the 9-1-1 gate, as indicated by the curved red arrow. The green Rad24 upper loop, with the help of the cyan Rfc5 plug, prevent DNA with a gap size shorter than 5 nt from entering the Rad24-RFC chamber. The consistent presence of four ATPγS and one ADP thorough out steps 1 to 4 suggests that ATP hydrolysis is not required for DNA binding and clamp gate closure. Stable DNA binding and clamp gate closure likely induce a conformational change in Rad24-RFC to trigger ATP hydrolysis, leading to the dissociation of Rad24-RFC from 9-1-1 in step 6, leaving 9-1-1 alone encircling the DNA 3′-end. **b**) A sketch illustrating that Rad24-RFC does not load 9-1-1 onto a nicked DNA or very short gap (1-4 nt). By virtue of its recognition to the 5′-end, Rad24-RFC loads 9-1-1 to the 3′-end of a medium sized gap but loading at a very long gap or a 5′-recessed DNA, leads to loading onto ssDNA at a 5’ ssDNA junction which is a very different outcome. Loading on the duplex of a 3’ end enables the 9-1-1 clamp to be used by a Pol, such as TLS Pol recruitment for gap filling at 3′ end, while loading at a 5’ end may result in ATM activation and cell cycle arrest.

### 3. Loading of 9-1-1 on 5′ sites by Rad24-RFC is much slower than PCNA loading by RFC

We demonstrate here that 9-1-1 clamp loading at a 10-nt DNA gap is much slower than RFC loading PCNA at the same gap (**Fig. 1b**). This finding is consistent with the different ATPase activity of human Rad17-RFC compared to human canonical RFC at a 5′ ss/ds DNA (39). Thus, when normal repair pathways of a cell are not “overloaded”, PCNA is likely assembled onto gapped DNA and utilized for repair rather than a 9-1-1 clamp that may be utilized to activate the DNA damage checkpoint and shut down the cell cycle. Presumably, in the event of significant DNA damage, the 9-1-1 clamp is loaded at stalled and incomplete Okazaki fragments or DNA repair gaps to signal the DNA damage response and to halt the cell cycle to gain time to repair DNA before continuing the cell cycle.

### 4. Possible physiological consequences of the different DNA structures used by Rad24-RFC and RFC

The replicative clamp loader RFC is well known to load PCNA at a 3′-recessed DNA end, the normal primer for DNA synthetic extension (32). In contrast, Rad24-RFC has been established to load the damage signaling 9-1-1 clamp onto a 5′-recessed DNA (39, 41). However, three recent structural studies on RFC have broken the established norm, revealing the 5′-recessed DNA binding capacity on the shoulder of the Rfc1 subunit of RFC and the loading of PCNA onto a gapped DNA (at the 3′ end) where both 3′ and 5′-recessed ends are present (28-30). The current study shows the Rad24-RFC, like RFC, is also able to load 9-1-1 onto a gapped DNA by simultaneously binding to both the 5′-recessed and 3′-recessed DNA. This raises the question of whether there is any significant mechanistic difference in loading their respective clamps onto gapped DNA substrates. Based on the current structural and biochemical information, we suggest two major differences (**Fig. 6b**). Firstly, Rad24-RFC can’t unwind dsDNA and thus requires a pre-existing ssDNA gap of a minimum length of 5-nt for efficient loading while RFC can unwind DNA and even load PCNA at nicks, due to its ability to unwind DNA at a nick or small DNA gaps (28-30). Secondly, Rad24-RFC may be evolved to prefer ssDNA gaps so as not to trigger a cell cycle checkpoint over nicks that can be sealed by ligase or dealt with by short gap base excision repair (i.e., a 1-nt gap).

In overview, eukaryotes have evolved different clamp loaders to efficiently maintain genome integrity. For many short ssDNA patches (< 5-nt gap) produced during base excision pathways, RFC is likely utilized to load PCNA to these sites, which goes on to recruit a translesion synthesis polymerase (e.g., Pol β, Pol ε, Pol ζ, and Rev1) to fill in the short gaps, followed by the action of a ligase to seal the nick and complete repair (54-56). Rad24-RFC seems to be reserved for ssDNA patches of ≥ 5-nt. This might enable eukaryotic cells to efficiently repair very short gap DNA damage, without triggering the cell cycle 9-1-1 activated ATR kinase route that might be more important when extensive damage and longer ssDNA gaps occur. However, it is interesting to note that 9-1-1 interacts with Pol ζ to reduce Pol ζ-dependent spontaneous mutagenesis (45) and with Pol ε to increase the catalytic efficiency for correct nucleotide incorporation (57). Furthermore, 9-1-1 interacts with TopBP1 and DNA polymerase-α directly at stalled DNA replication forks (58) and with Pol β to stimulate its activity (46). Therefore, 9-1-1 likely has a broader role that functions beyond DNA damage signaling. The ability of Rad24-RFC to load 9-1-1 clamps at 5-10 nt gaps indicates that the 911 clamp will be left (after clamp loader ejection) to encircle the 3′ duplex, not the 5′ duplex, of the medium-sized gaps, and this places its orientation on DNA to favor action with DNA polymerases to fill-in gaps produced by DNA damage.

#### Limitations of the study

The current study has revealed a multi-step loading mode of the 9-1-1 clamp onto – unexpectedly – the 3′ end DNA of a medium sized gap (5-30 nt) by the alternative clamp loader Rad24-RFC. This discovery has enabled us to propose a new function for 9-1-1 in DNA damage repair, specifically that this process may enable the use of the 9-1-1 clamp by DNA polymerases for gap repair. The proposed function needs further in vivo studies. However, a more detailed understanding also requires more extensive biochemical, biophysical and structural investigation.

## STAR ⋆ METHODS

Detailed methods are provided in the online version of this paper and include the following:

- KEY RESOURCES TABLE
- RESOURCE AVAILABILITY
  - Lead contact
  - Materials availability
  - Data and code availability
- EXPERIMENTAL MODEL AND SUBJECT DETAILS
- METHOD DETAILS
  - Protein expression and purification
  - Magnetic bead based 9-1-1 clamp loading assays
  - Cryo-EM grids preparation and data collection
  - Image processing and 3D reconstruction
  - Model building, refinement, and validation
- QUANTIFICATION AND STATISTICAL ANALYSIS

## SUPPLEMENTAL INFORMATION

Supplemental Information can be found online at:

## ACKNOWLEDGEMENTS

Cryo-EM micrographs were collected at the David Van Andel Advanced Cryo-Electron Microscopy Suite in Van Andel Institute. We thank G. Zhao and X. Meng for facilitating data collection. This work was supported by the US National Institutes of Health grants GM131754 (to H.L.) and GM115809 (to M.E.O.), Van Andel Institute (to H.L.), and Howard Hughes Medical Institute and BCRF 22-068 (to M.E.O.).

## AUTHOR CONTRIBUTIONS

M.E.O. and H.L. designed research; F.Z., R.E.G. and N.Y.Y. performed research; F.Z., R.E.G., N.Y.Y., M.E.O., and H.L. analyzed the data; F.Z., M.E.O., and H.L. wrote the manuscript with input from all authors.

## STAR ⋆ METHODS

## RESOURCE AVAILABILITY

### Lead contact

Further information and requests for reagents and resources may be directed to, and will be fulfilled by the Lead Contact Huilin Li

### Materials availability

All plasmids used in this study will be available from the lead contact upon request. This study did not generate new unique reagents.

### Data and code availability

- The five EM maps of the *S. cerevisiae* Rad24-RFC–9-1-1 clamp–10-nt gapped DNA complex (steps 1-5) and one EM map of the Rad24-RFC–9-1-1 clamp–5-nt gapped DNA complex have been deposited in the Electron Microscopy Data Bank with accession codes EMD-29412, EMD-29413, EMD-29414, EMD-29415, EMD-29416 and EMD-29417 respectively. Their corresponding atomic models have been deposited in the Protein Data Bank with accession codes 8FS3, 8FS4, 8FS5, 8FS6, 8FS7, and 8FS8. These data will be publicly available as of the date of publication.
- This paper does not report original code.
- Any additional information required to reanalyze the data reported in this work will be available from the lead contact upon request.

## EXPERIMENTAL MODEL AND SUBJECT DETAILS

The expression plasmids of *S.c.* Rad24-RFC, 9-1-1 clamp and RPA were generated by standard molecular biology techniques, constructed using the DH5α strain of *E. coli* (Thermo Fisher Scientific), and are listed in the key resources table. Those three multiprotein complexes were purified from the BL21(DE3) strain of *E. coli* (Thermo Fisher Scientific). *E. coli* strains were cultured in LB media.

## METHOD DETAILS

### Protein expression and purification

Rad24-RFC and the 9-1-1 clamp were expressed and purified as described in our earlier report (43). RPA was purified as described in (59).

### Magnetic bead-based 9-1-1 clamp loading assays

S.c. 9-1-1 having an N-terminal 7-residue kinase recognition sequence on Mec3 was radiolabeled to a specific activity of approximately 70 cpm/fmol with [γ-^32^P] ATP (PerkinElmer Life Sciences, Inc) using the recombinant catalytic subunit of cAMP-dependent protein kinase produced in *E. coli* (a gift from Dr. Susan Taylor, University of California at San Diego) as described (60). Protein concentrations were determined by Bradford assay reagent (Bio-Rad) using BSA as a standard. The oligonucleotides used to make the DNA substrates in this report (**Supplementary Table 1**) were synthesized and PAGE purified or HPLC purified by Integrated DNA Technologies. To form the isolated nicked and gapped templates, 2500 pmol of each of primer #1 and primer #2 were mixed with 1250 pmol of the template strand in 45 μl of Buffer A (5 mM Tris-HCl, 150 mM NaCl, 15 mM sodium citrate, final pH 8.5), then incubated in a 95°C water bath and cooled to room temperature (23°C) over a 30-min interval. 200 pmol of the annealed primed templates were combined with 1 mg of Dynabeads M-280 Streptavidin beads (ThermoFisher Scientific) in 200 μl of Buffer B (40 mM HEPES-NaOH pH 7.5, 0.1 mg/ml BSA, 1 mM DTT, 8 mM MgCl_2_ and 120 mM NaCl) and incubated at 23 °C for 30 minutes with agitation. Supernatant containing excess unbound primer oligonucleotides was removed using a magnetic separator. The DNA-beads were washed twice with 250 μl Buffer C (30 mM HEPES-NaOH pH 7.5, 1 mM DTT, 1 mM CHAPS) with 7 mM MgCl_2_ and 100 mM NaCl, and then resuspended in 100 μl of the same buffer. The concentration of DNA conjugated to the beads was determined by measuring absorbance at 260 nm using a nanodrop spectrophotometer and was typically in the range of 1.0 to 2.0 pmol DNA per μl bead mix.

Before the start of the 9-1-1 loading assay, the Dig at the end of the DNA-bead conjugate was first blocked with anti-digoxigenin Fab fragment (Roche) in a ratio of 1:2 respectively in Buffer C with 7 mM MgCl_2_. After the mixture was incubated at room temperature for 15 minutes with agitation, RPA was added to the DNA-bead conjugate in a ratio of 4:1 and incubated at 30°C for 5 minutes. The supernatant containing excess unbound Fab fragments and RPA was then removed using a magnetic separator. The blocked DNA-beads were resuspended in the following clamp loading mixture. Each reaction contained 100 nM antibody-blocked DNA-bead conjugate, 200 nM ^32^P-9-1-1 in 50 μl of Buffer C with 8 mM MgCl_2_, 50 mM NaCl and 1 mM ATP. The clamp loading reaction was initiated with 174 nM Rad24-RFC and incubated at 30°C with agitation for 5 minutes. The reaction was stopped by quenching with 25 mM EDTA on ice. The beads were then isolated with a magnetic separator and free protein was washed away with two washes of 250 μl Buffer C containing 100 mM NaCl. The DNA-bound radioactive 9-1-1 was stripped from the beads with 0.5% SDS and 5 minutes of boiling. One half (25 μl) of the supernatant was then counted by liquid scintillation and quantitation was multiplied by two for quantitation of the full reaction.

For the time course comparison of PCNA loading versus 9-1-1 loading, the 10-nt gapped DNA was used and each reaction contained 100 nM antibody-blocked DNA-bead conjugate, 200 nM ^32^P-PCNA or 200 nM ^32^P-9-1-1 and 150 nM RFC or 150 nM Rad24-RFC in 50 μl of Buffer C with 8 mM MgCl_2_ and 1 mM ATP. The PCNA loading reactions were incubated on ice for 0 to 25 seconds before being quenched with 25 mM EDTA. The 9-1-1 loading reactions were also incubated on ice but for longer times (0 to 4 minutes) before being quenched with EDTA. The amount of radiolabeled clamp loaded in each reaction was determined as described above.

### Cryo-EM grids preparation and data collection

The 10-nt gapped DNA substrate with both 3′- and 5′-junctions, and its sequence is shown in **Fig. 2a**. This DNA is the same as used in the biochemical assays as described above and was prepared as described previously (29). The in vitro assembly of yeast Rad24-RFC–9-1-1 clamp–10-nt gapped DNA complex followed our previous procedure for reconstituting yeast Pol δ–PCNA–DNA complex (61). Briefly, 6.6 μl purified 9-1-1 protein at 5.0 μM and 3.7 μl gapped DNA at 11.2 μM were mixed and incubated at 30°C for 10 min, then the mixtures, along with 0.75 μl 10 mM ATPγS and 0.75 μl 100 mM Mg-Acetate, were each added into 3.2 μl purified Rad24-RFC protein at 8.6 μM concentration, the final concentrations of the components were: Rad24-RFC at 1.8 μM, 9-1-1 clamp at 2.2 μM, 10-nt gapped DNA at 2.8 μM, ATPγS at 0.5 mM, and Mg-Acetate at 5 mM; the total reaction volume was 15 μl. The final molar ratio of Rad24-RFC: 9-1-1 clamp: DNA was 1.0: 1.2: 1.6. The mixture was then incubated in an ice-water bath for 1.5 hr. Assembly of Rad24-RFC–9-1-1 clamp with the 5-nt gapped DNA (**Fig. 5a**) followed the same procedure as the 10-nt gapped DNA, except that the gapped region is shortened to 5 (polydT_5_).

The Quantifoil Cu R2/1 300 mesh grids were glow discharged for 1 min in a Gatan Solarus, then 3 μl of the mixture was applied onto the EM grids. Sample vitrification was carried out in a Vitrobot (Thermo Fisher Mark IV) with the following settings: blot time 2.5 s, blot force 3, wait time 3 s, inner chamber temperature 6°C with 95% relative humidity. The EM grids were flash-frozen in liquid ethane cooled by liquid nitrogen. Cryo-EM data were automatically collected on a 300 kV Titian Krios electron microscope controlled by SerialEM (62) in a multi-hole mode. The micrographs were captured at a scope magnification of 105,000×, with the objective lens under-focus values ranging from 1.3 to 1.9 μm, by a K3 direct electron detector (Gatan) operated in the super-resolution video mode. During a 1.5 s exposure time, a total of 75 frames were recorded with a total dose of 64 e^-^/Å^2^. The calibrated physical pixel size was 0.828 Å for all digital micrographs.

### Image processing and 3D reconstruction

The data collection and image quality were monitored by the cryoSPARC Live v3.1 (50) installed in a local workstation. The image preprocessing including patch motion correction (on bin x2 data), contrast transfer function (CTF) estimation and correction, blob particle picking (70-150 Å diameter), and particle extraction (on bin x4 data) were also achieved at the same time. A total of 19,603 raw micrographs for the 10-nt gapped DNA bound Rad24–RFC-911 complex was recorded during a three-day data collecting session (**Supplementary Fig. 2**). The extracted particle images were subjected to two rounds of 2-dimensional (2D) classification, resulting in a selected dataset of ∼2.4 million “good” particle images. We also trained the automatic particle picking program Topaz (63) and used the trained model to pick up another particle dataset. The Topaz picked dataset was also subjected to two rounds of 2D classifications, resulting in a selected dataset of ∼2.9 million “good” particle images. We then used the reported “Build and Retrieve” method to retrieve those less frequently occurring particle views (64). The two particle datasets were combined by removing duplicates that were 40% overlapping (52 Å) or larger than the preset particle diameter (∼130 Å). This resulted in a merged dataset of ∼3.3 million particle images.

We performed ab initio 3D reconstruction with the combined dataset and obtained three initial maps in cryoSPARC. One map was chosen as reference for later 3D classification. The particle coordinates were transformed into Relion format (51) using the PyEM program (65). At the same time, the collected raw images are imported into Relion 4.0, motion corrected by MotionCor2 (66), and CTF estimated and corrected by CTFFIND4 (67). Then, the particle images were re-extracted with the particle coordinates imported from cryoSPARC. We next performed 3D classification in Relion and obtained six 3D classes. One map lacked feature and was considered “junk”, but the remaining 5 maps corresponded to different stages of 9-1-1 clamp loading by Rad24-RFC onto the 10-nt gapped DNA (**Supplementary Fig. 2**). After 3D auto-refinement, CTF refinement and Bayesian polish in Relion, the particles from each class were imported back to cryoSPARC for final non-uniform refinement, which resulted in five maps of the Rad24-RFC–9-1-1–10-nt gapped DNA complex. The map with an open 9-1-1 ring but lacking DNA density in the central chamber of Rad24-RFC was at 2.93 Å, the map with an open 9-1-1 ring and a weak chamber DNA density was at 2.94 Å, the map with an open 9-1-1 ring and a strong chamber DNA density at 2.76 Å, the map with a partially closed 9-1-1 ring and a strong chamber DNA density was at 2.90 Å, and the map with a fully closed 9-1-1 ring and a strong chamber DNA density was at 2.85 Å average resolution. All five maps had a 5′ DNA stably bound at the Rad24 external shoulder site (**Supplementary Figs 2-3**).

A similar data collection and image processing strategy was used for the 5-nt gapped DNA bound Rad24-RFC–9-1-1 complex **(Supplementary Fig. 8**). We collected a dataset of 22,570 raw micrographs. The initial 3D reconstruction and heterogeneous refinement resulted in only two conformations, different from the five conformations of the 10-nt gapped DNA bound complex. One map was constructed from ∼3.5 million particles and had an open 9-1-1 ring. This map had clear density for the 5′ shoulder DNA but had neither DNA density in the open 9-1-1 ring nor in the Rad24-RFC chamber. This map was essentially the same as the step-1 map we have solved for the 10-nt gapped DNA bound complex, thus was not processed further. The other map was reconstructed from about 2.1 million particles and had a closed 9-1-1 ring. This map had weak DNA density inside the Rad24-RFC chamber. To improve the chamber DNA density, we first performed 3D classification in cryoSPARC (50) and obtained 20 3D classes. Among the 20 maps, four had strong DNA density in the central chamber of Rad24-RFC and they were combined and imported into Relion (51) for further focused 3D classification around the chamber containing DNA region. We finally isolated a subpopulation of 242,906 particles with a relative stable chamber DNA density for final refinement, leading to the presented EM map at 3.04 Å average resolution (**Supplementary Figs. 3f-g and 8**).

### Model building, refinement, and validation

The previously reported cryo-EM structure of the yeast Rad24-RFC–9-1-1 clamp complexed with a shoulder DNA (PDB entry 7SGZ) was used as the starting model for atomic modeling of the six EM maps (52). To model DNA in the central chamber of the clamp loader, we started with the 3′ ss/ds DNA-central chamber coordinates extracted from the structure of the yeast RFC–PCNA–2 DNA complex structure (PDB entry 7TFH), followed by manual adjustment. We also used the de novo modeling program Map-to-Model wrapped in PHENIX (68) and the module “Automated Nucleic Acid building” function integrated in COOT (69). The two approaches led to similar gapped DNA models. The initial DNA model was first fitted into the most robust step-5 3D map of the 10-nt gapped DNA bound complex with the closed 9-1-1 ring, then merged as a single coordinate file in UCSF Chimera (70). This served as the starting model for the step-5 EM map. The model was refined iteratively between the real space refinement in PHENIX and the manual adjustment in COOT. The atomic model of the step-5 complex was refined to 2.9 Å, and was comprehensively validated by the MolProbity program (71) embedded in PHENIX. This model then served as the initial model for the four other 10-nt gapped DNA bound maps and the single 5-nt gapped DNA bound complex map. The model for each 3D map was subjected to a similar manual building process (**Supplementary Table 2**). The step-1 and -2 maps with 10-nt gapped DNA had weaker EM densities for the two 9-1-1 gate subunits Ddc1 and Mec3, and the map of the 5-nt gapped DNA bound complex had weak 3′ ss/ds DNA density in the chamber. We used rigid body docking in these weak density regions, followed by manual adjustment in COOT and refinement by PHENIX. Structure figures were prepared using ChimeraX (72), and assembled and labelled in Adobe Illustrator (Adobe Inc, San Jose, CA).

## QUANTIFICATION AND STATISTICAL ANALYSIS

Statistical analyses were performed using GraphPad Prism (GraphPad Software Inc.). Data are expressed as the mean ± SD.

## Supplementary Information

**Supplementary Table 1.**
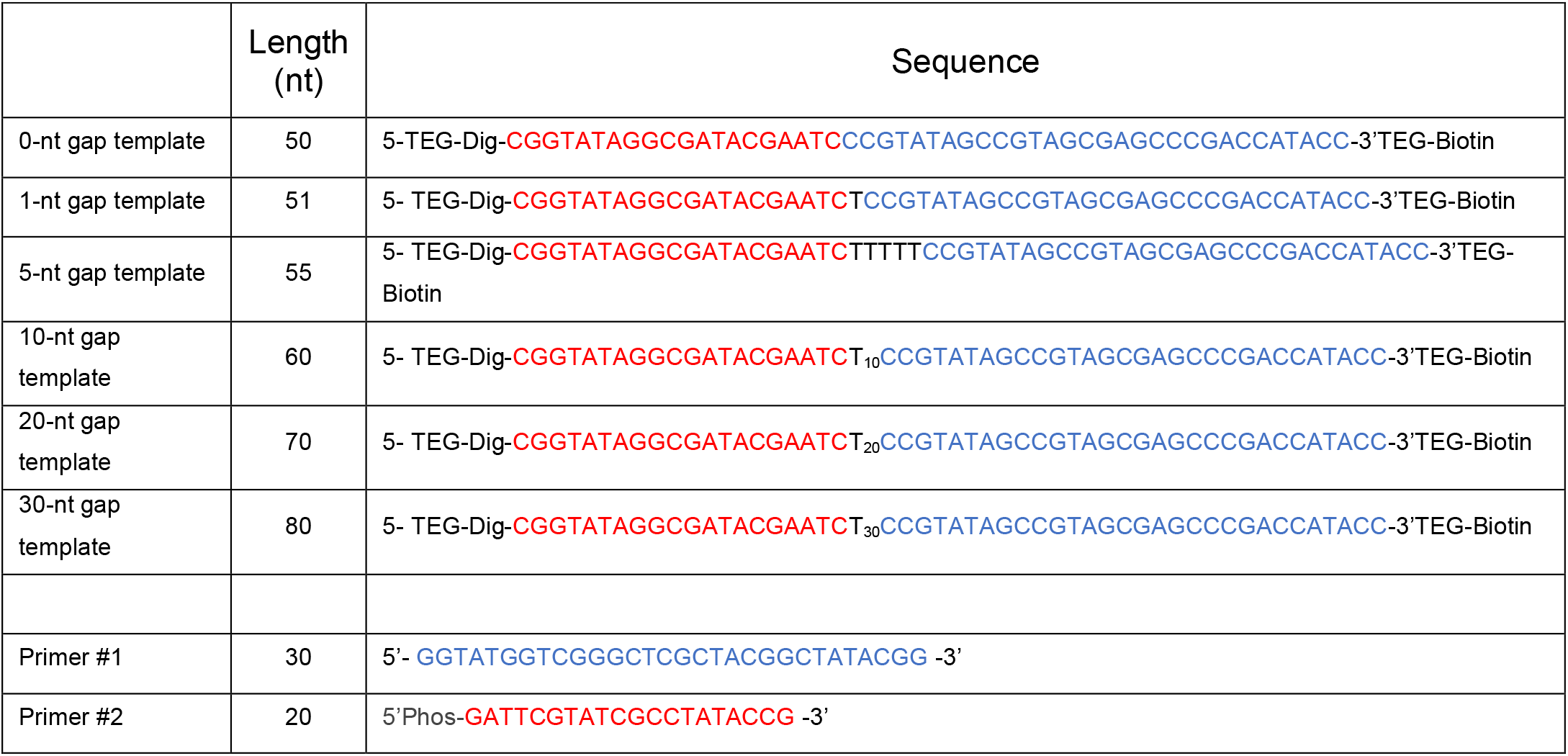
Oligonucleotides used for in vitro 9-1-1 loading experiments. Template nucleotides in red type are complimentary to the Primer #2 sequence, and template residues in blue type are complimentary to the Primer #1 sequence. Template nucleotides in black type are the ssDNA gap.

**Supplementary Table 2.** Cryo-EM data collection, refinement, and atomic model validation of the 10-nt and the 5-nt gapped DNA-bound yeast Rad24-RFC−9-1-1 complexes.

| Structures | Rad24-RFC–9-1-1–10-nt gapped DNA |  |  |  |  | 5-nt gapped DNA bound |
| --- | --- | --- | --- | --- | --- | --- |
|  | Step 1 | Step 2 | Step 3 | Step 4 | Step 5 |  |
| Data collection and processing |  |  |  |  |  |  |
| Magnification |  |  | 105,000 |  |  |  |
| Voltage (kV) |  |  | 300 |  |  |  |
| Electron dose (e-/Å²) |  |  | 64 |  |  |  |
| Under-focus range (µm) |  |  | 1.3 - 1.9 |  |  |  |
| Pixel size (Å) |  |  | 0.828 |  |  |  |
| Symmetry imposed |  |  | C1 |  |  |  |
| Initial particle images (no.) |  |  | 3,256,498 |  |  | 5,857,353 |
| Final particle images (no.) | 255,777 | 347,098 | 400,932 | 191,448 | 292,725 | 242,906 |
| Map resolution (Å) | 2.93 | 2.94 | 2.76 | 2.90 | 2.85 | 3.04 |
| FSC threshold |  |  |  | 0.143 |  |  |
| Map resolution range (Å) | 2.0 – 12.0 | 2.0 – 11.0 | 2.0 – 12.0 | 2.0 – 12.0 | 2.0 – 11.0 | 2.0 – 13.0 |
| Refinement |  |  |  |  |  |  |
| Initial model used (PDB code) |  | Step-5 model of this study |  |  | 7SGZ, 7TFH | Step-5 |
| Map sharpening B factor (Å²) | -86.9 | -76.6 | -82.6 | -77.6 | -82.0 | -72.4 |
| Map to model CC<sub>mask</sub> | 0.81 | 0.78 | 0.81 | 0.81 | 0.84 | 0.75 |
| Model composition |  |  |  |  |  |  |
| Non-hydrogen atoms | 21,221 | 21,450 | 22,206 | 22,260 | 22,190 | 21,901 |
| Protein and DNA residues | 2,571; 27 | 2,538; 50 | 2,618; 58 | 2,623; 58 | 2,615; 58 | 2,581; 55 |
| Ligands | 9 | 9 | 9 | 9 | 9 | 9 |
| R.m.s. deviations |  |  |  |  |  |  |
| Bond lengths (Å) | 0.004 | 0.004 | 0.003 | 0.003 | 0.003 | 0.002 |
| Bond angels (°) | 0.642 | 0.632 | 0.680 | 0.648 | 0.555 | 0.511 |
| Validation |  |  |  |  |  |  |
| MolProbity score | 2.32 | 2.07 | 1.82 | 1.97 | 1.85 | 2.04 |
| Clashscore | 11.23 | 9.63 | 7.23 | 7.86 | 6.64 | 9.09 |
| Poor rotamers (%) | 4.18 | 2.65 | 1.73 | 2.53 | 2.28 | 3.17 |
| Ramachandran plot |  |  |  |  |  |  |
| Favored (%) | 95.90 | 96.35 | 96.38 | 96.43 | 96.65 | 96.99 |
| Allowed (%) | 4.10 | 3.65 | 3.62 | 3.57 | 3.35 | 3.01 |
| Disallowed (%) | 0 | 0 | 0 | 0 | 0 | 0 |

**Supplementary Fig. 1.**
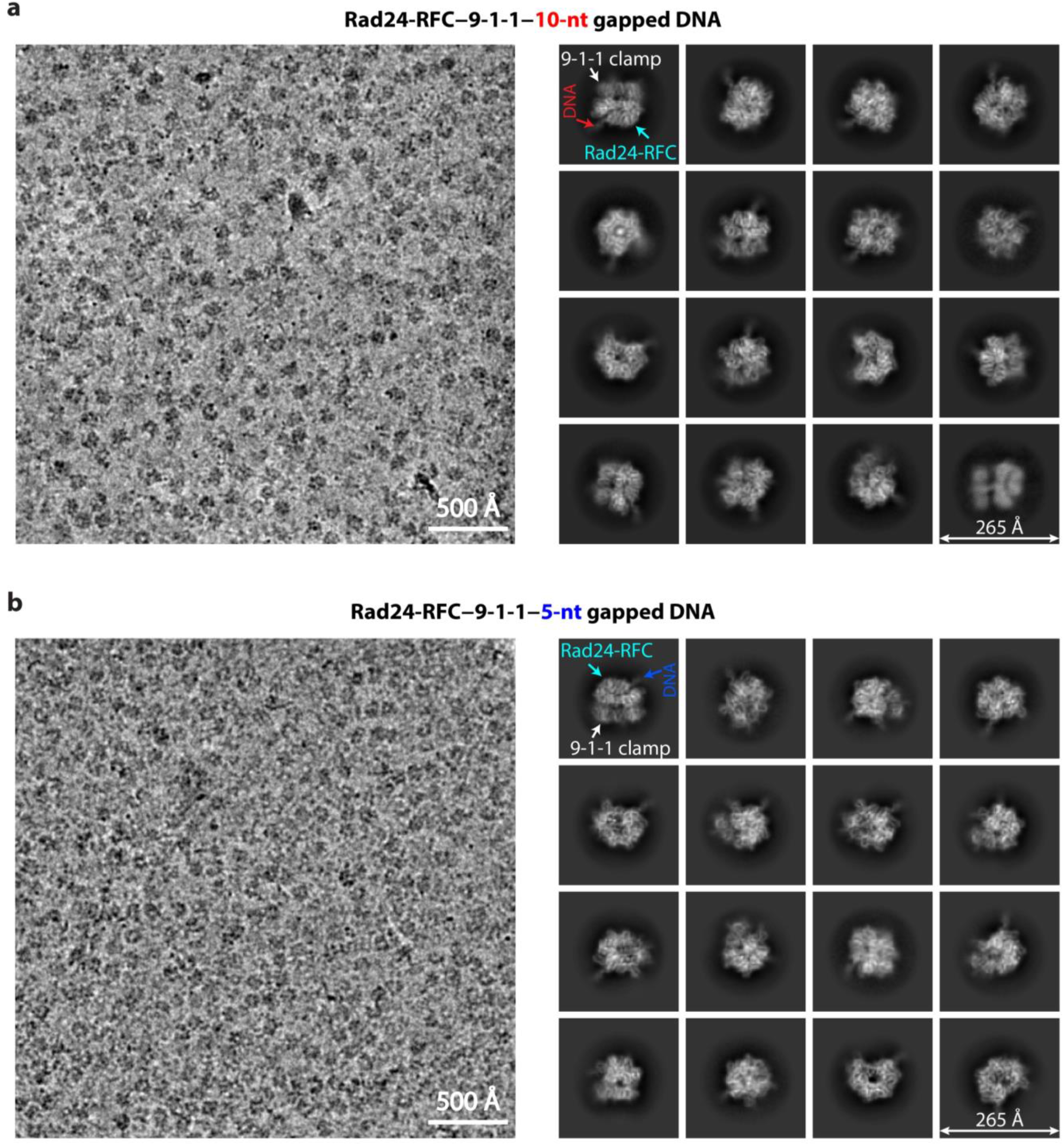
Cryo-EM of S. cerevisiae Rad24-RFC–9-1-1 clamp–gapped DNA complexes. **a-b**) Left panels show a typical raw micrograph of the Rad24-RFC–9-1-1 clamp bound to a 10-nt (a) or a 5-nt (b) gapped DNA assembled in vitro by directly mixing separately purified components. Right panels show selected 2D class averages in different views. The EM densities for Rad24-RFC, 9-1-1, and DNA are labelled in the first 2D averages. Scale bar is shown in each panel.

**Supplementary Fig. 2.**
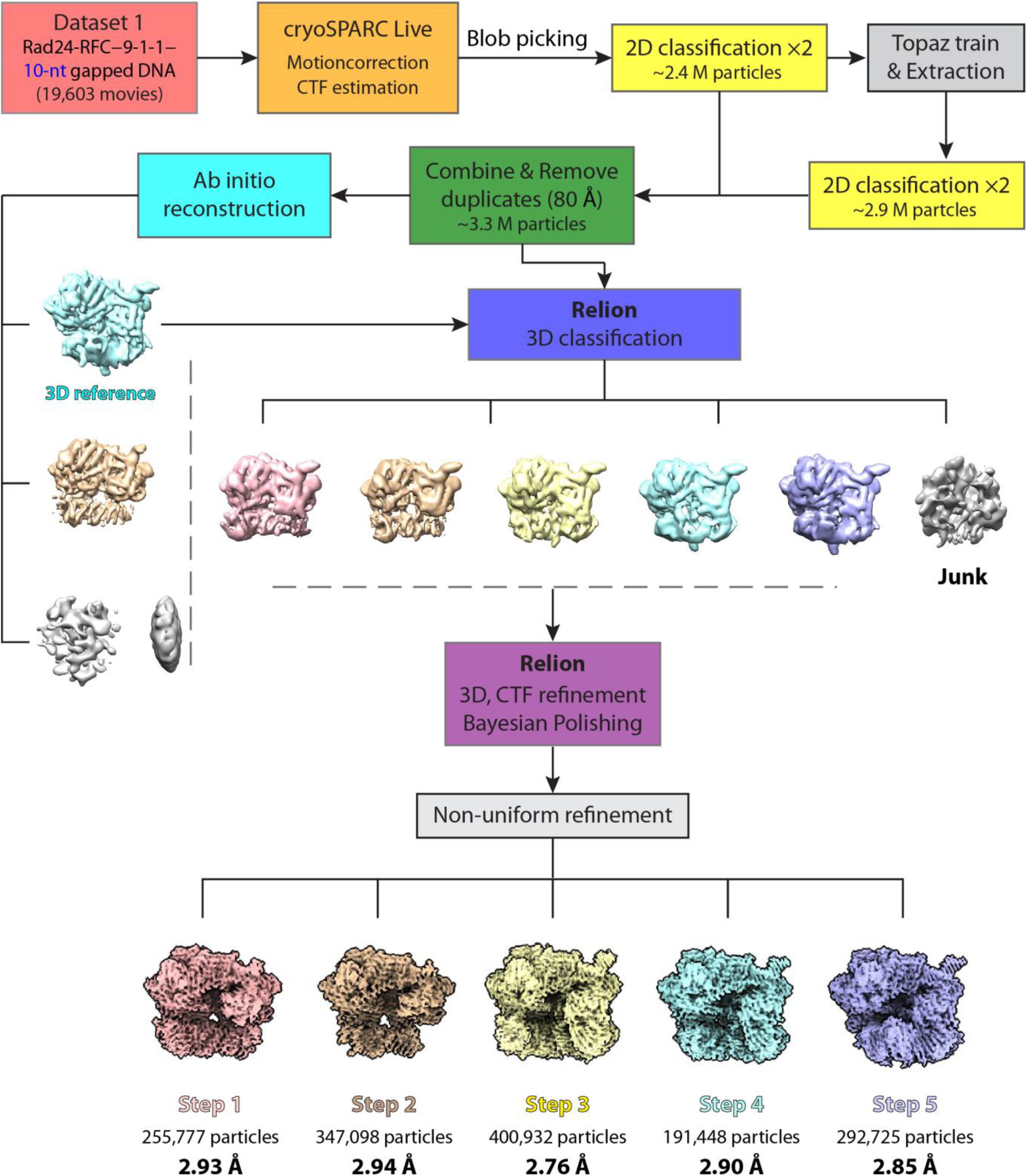
Workflow of cryo-EM data processing and 3D reconstruction of the 10-nt gapped DNA bound Rad24-RFC–9-1-1 clamp complex. Approximately 20,000 movie micrographs were collected. Data collection was monitored by cryoSPARC Live (v3.2), which was also used for pre-processing, particle picking, and initial 3D reconstruction. Next, the particle images and produced initial 3D reference map were imported into Relion (v4.0 beta) for 3D classification, CTF refinement, and Bayesian polishing. Finally, the data were imported back to cryoSPARC for non-uniform refinement, leading to the five intermediate structures. See **Methods** for more details.

**Supplementary Fig. 3.**
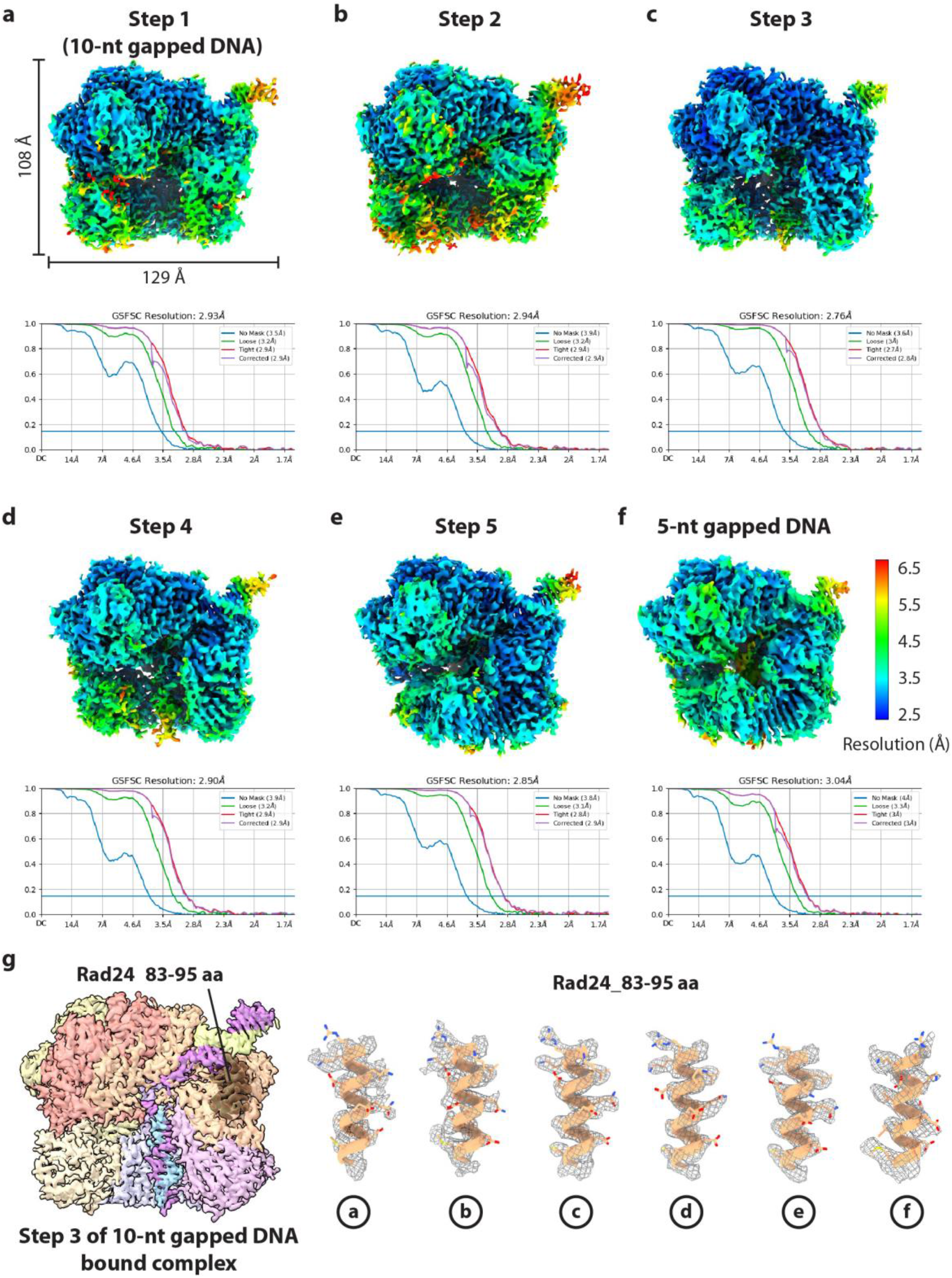
Local resolution maps, resolution estimation, and local density features of the six EM maps of the 10-nt or 5-nt gapped DNA bound complexes. **a-f**) Upper: Color-coded local resolution maps of the five 10-nt (a-e), and one 5-nt (f) gapped DNA bound EM maps. Lower: Gold standard Fourier shell correlation curves (GSFSC) of each map. The size of all maps is similar and labeled only on the first map. **g**) Left: surface-rendered EM map of 10-nt gapped DNA bound complex in step 3, highlighting an α-helix (aa 83-95) of Rad24 in dark orange. This helix in each of the six maps is enlarged and shown in the right panel, with the EM density rendered in grey meshes, atomic model in cartoons, and side chains in sticks.

**Supplementary Fig. 4.**
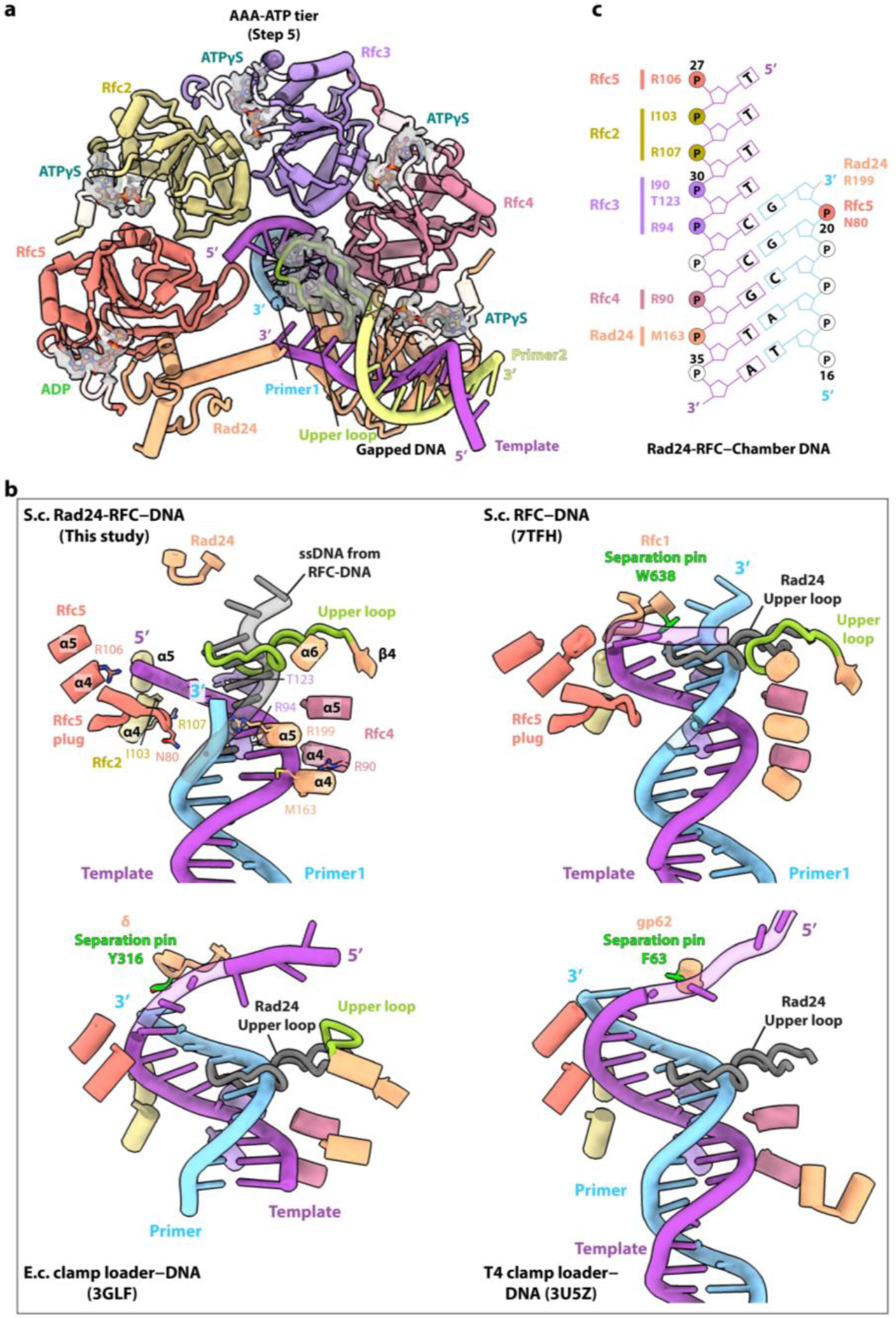
Rad24-RFC possesses a unique long loop that blocks 3′ chamber DNA from moving upwards. **a)** EM densities of the bound nucleotides and the Rad24 upper long loop in the 10-nt gapped DNA-bound step-5 structure. The structure is showed here as an example, as four ATPγS and one ADP are resolved in all six EM maps (five 10-nt gapped DNA bound and one 5-nt gapped DNA bound) captured in this study. Only the AAA-ATP tier is shown in top cartoon view for clarity. The EM map within 6.5 Å around the nucleotides and the Rad24 upper loop were rendered at a threshold of 0.14 and shown as transparent grey surfaces. The five nucleotides are shown in sticks, and the Rad24 upper long loop is shown in green cartoon. **b)** Superposition of equivalent DNA binding loops in the chambers of different clamp–loader– DNA complexes. In the upper left panel Rad24-RFC-DNA is superimposed with the RFC-bound DNA (grey), and in remaining panels, the Rad24 upper long loop is shown in grey and superimposed to other loader-DNA structures as labeled. **c)** Sketch detailing Rad24-RFC residues interacting with the 3’ ss’ds DNA in the central chamber of Rad24-RFC. See also panel b and Fig. 3b.

**Supplementary Fig. 5.**
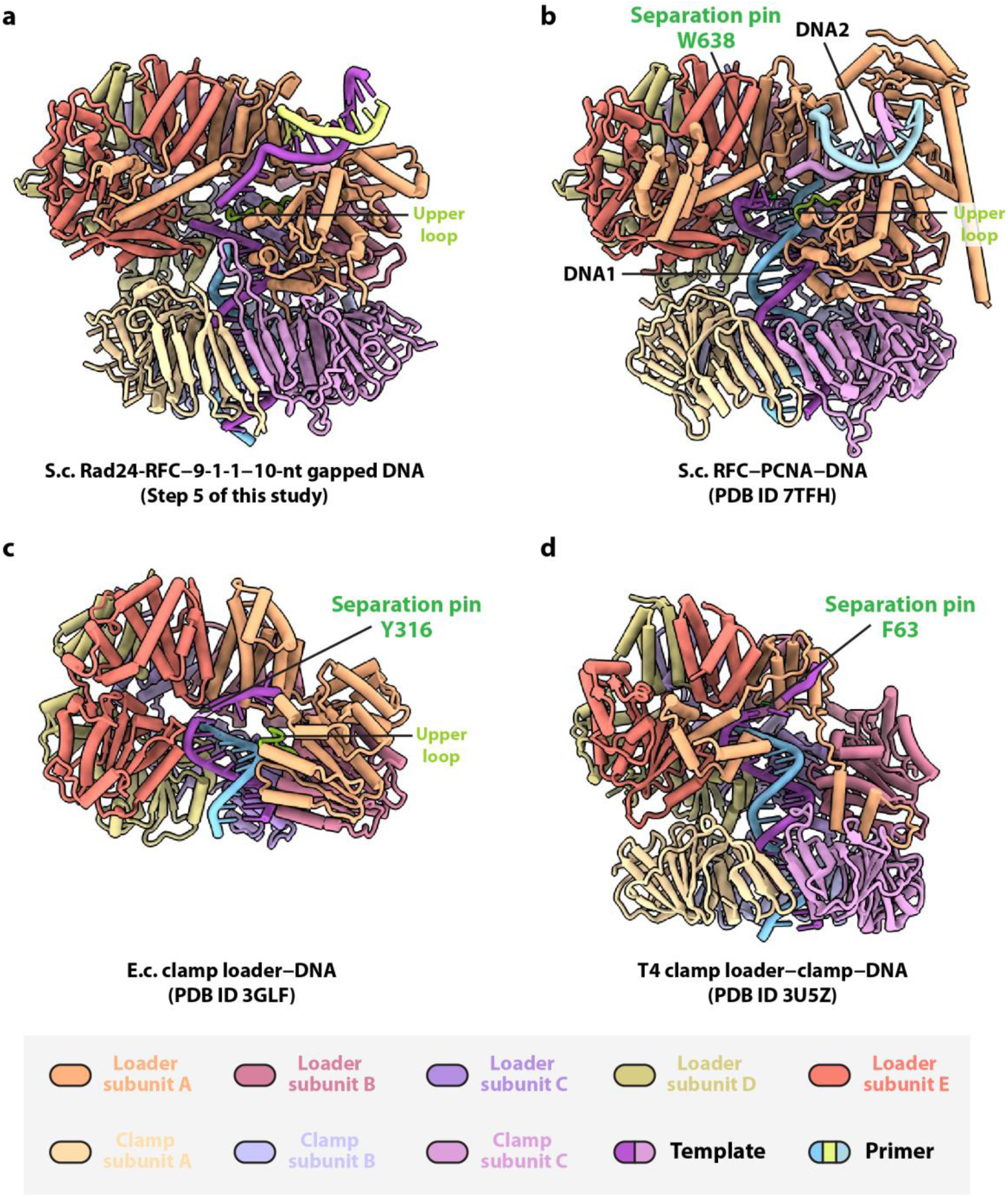
Comparison of clamp loader–[clamp]–DNA complex structures from different organisms reveals that the upper loop is uniquely long in Rad24. **a)** The step 5 structure of Rad24-RFC– 9-1-1 bound to 10-nt gapped DNA. **b)** Structure of two DNA bound yeast RFC−PCNA complex. **c)** Structure of the *E. coli* clamp loader–clamp–DNA complex. **d)** Structure of the T4 phage clamp loader–clamp complex. Note that T4 phage loader lacks a loop equivalent to the Rad24 upper loop, because its AAA+ module of subunit A is highly degenerated, retaining only two α-helices. These structures are aligned by the clamp loaders. The upper loop of loader subunit A is in green (also see Fig. 4b-e), and the separation pin residue right above the DNA 3′-junction – if present – is shown as lime sticks. The color scheme is shown at the bottom.

**Supplementary Fig. 6.**
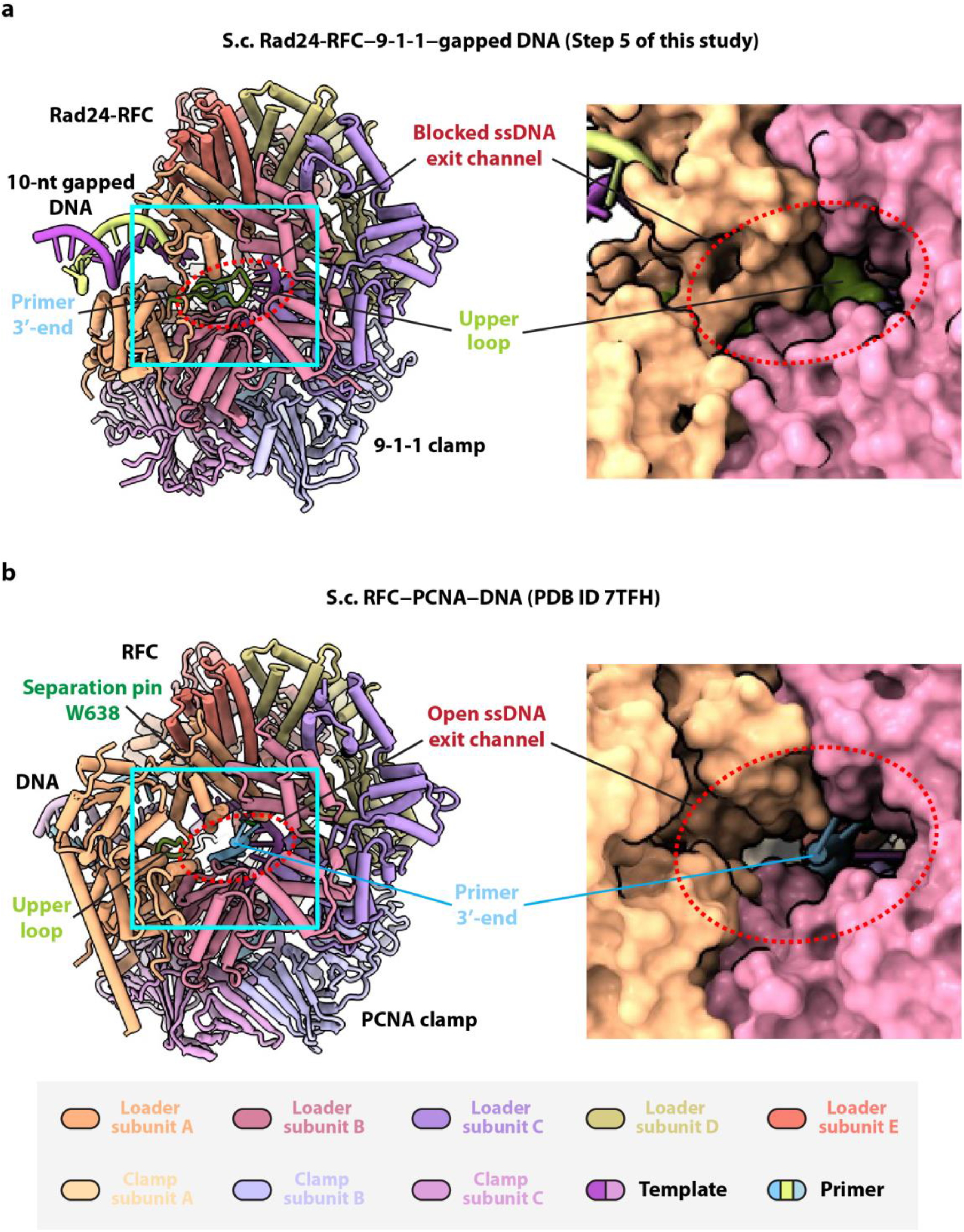
The 3′ ssDNA exit tunnel in RFC is blocked in Rad24-RFC. **a-b**) Left: Tilted views of Rad24-RFC−9-1-1 bound to 10-nt gapped DNA in step 5 (a) and RFC−PCNA bound to two DNA molecules (b). Right panels show the enlarged views of the 3′-junction regions marked by the cyan squares in the left panels. The dashed red oval marks the potential ssDNA exit tunnel in each panel. In Rad24-RFC, the exit tunnel of the ssDNA in the gap region is narrower and is fully blocked by the Rad24 upper loop (green). In contrast, the ssDNA exit tunnel is unblocked in RFC and in fact, the 2-nt ssDNA (cyan) - unwound by the Rfc1 separation pin (Trp-638) - is leaving the central chamber and emerging from the exit tunnel. The tunnel is formed by the loader subunits A and B and between the collar domains and AAA+ modules. Therefore, the primer 3′ ssDNA is prevented by the Rad24 upper loop from moving upwards to exit the chamber via the tunnel.

**Supplementary Fig. 7.**
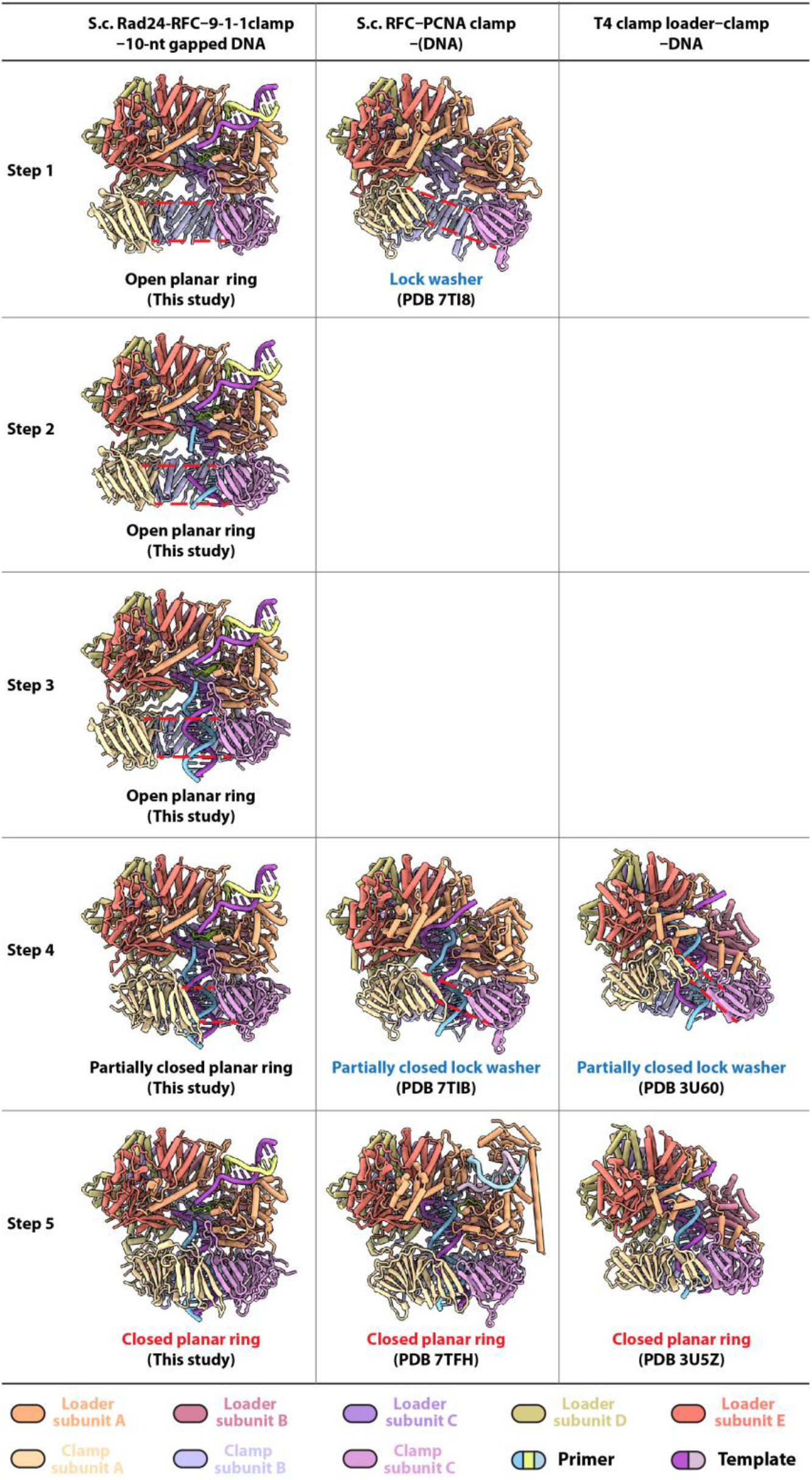
The planar open-gate 9-1-1 clamp versus the lock-washer shaped open-gate replication clamps (PCNA and T4 clamp). Column 1 shows the five intermediates of the yeast Rad24-RFC– 9-1-1 complex bound to 10-nt gapped DNA. The open or partially open-gate 9-1-1 clamps in steps 1 to 4 are all planar. Columns 2 and 3 show available clamp loader-clamp complexes in matching states: the yeast RFC– PCNA–DNA complexes (column 2), and the T4 phage clamp loader–clamp–DNA complexes (column 3). The replicative clamps are invariantly in a spiral or lock washer shape when in open-gate states. However, all clamps are planar when they are closed and encircle DNA. The parallel dashed red lines indicate the upper and lower bounds of the open DNA gate.

**Supplementary Fig. 8.**
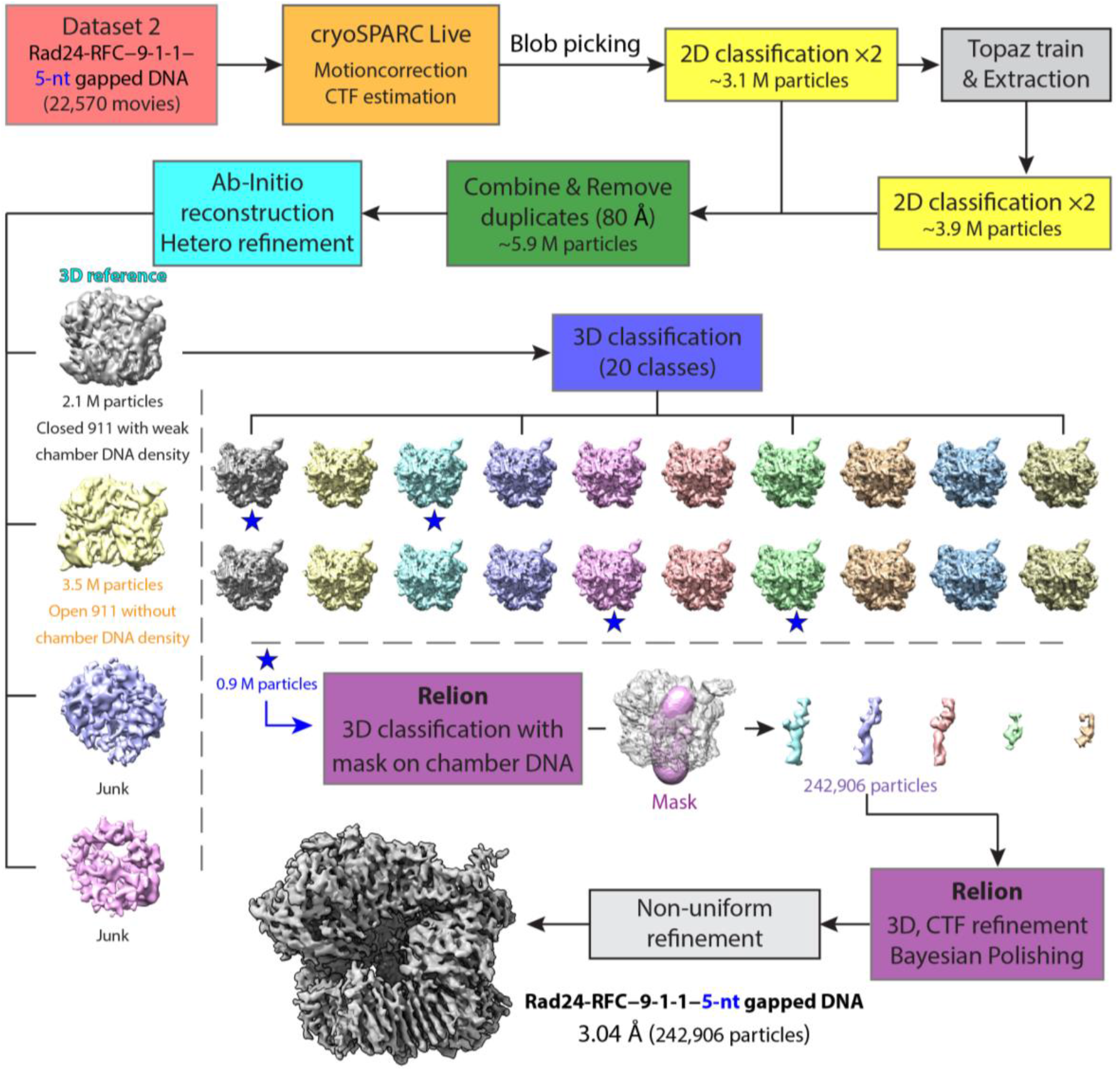
Workflow of cryo-EM data processing and 3D reconstruction of the 5-nt gapped DNA bound Rad24-RFC–9-1-1 complex. Approximately 23,000 raw video micrographs were collected. Data collection was monitored by cryoSPARC Live (v4.0), which was also used for pre-processing, particle picking, and initial 3D reconstruction, as described for the 10-nt gapped DNA bound dataset. After *ab initio* 3D reconstruction and heterogeneous refinement, four 3D classes were obtained. Two classes (purple and pink) were labeled “junk” as they lacked structural features, one map (yellow) had an open 9-1-1 clamp but no DNA density inside the chamber. The 4th 3D class, the grey map with 2.1 M (million) particles, had a closed 9-1-1 ring and weak density for DNA within the central chamber of the clamp loader; this class was chosen for further 3D classification leading to 20 3D classes. Four such classes labelled with blue stars had stronger DNA density in the central chamber, and they were combined (0.9 M particles) and imported into Relion (v4.0 beta) for further focused 3D classification with a mask (pink) around the chamber DNA. Among the five resulting 3D maps, the DNA density was distorted in four classes. The class with undistorted and strong DNA density (blue map with 242,906 particles) was imported back to cryoSPARC for final non-uniform refinement, leading to the final reported 3D map at 3.04 Å average resolution.

**Supplementary Fig. 9.**
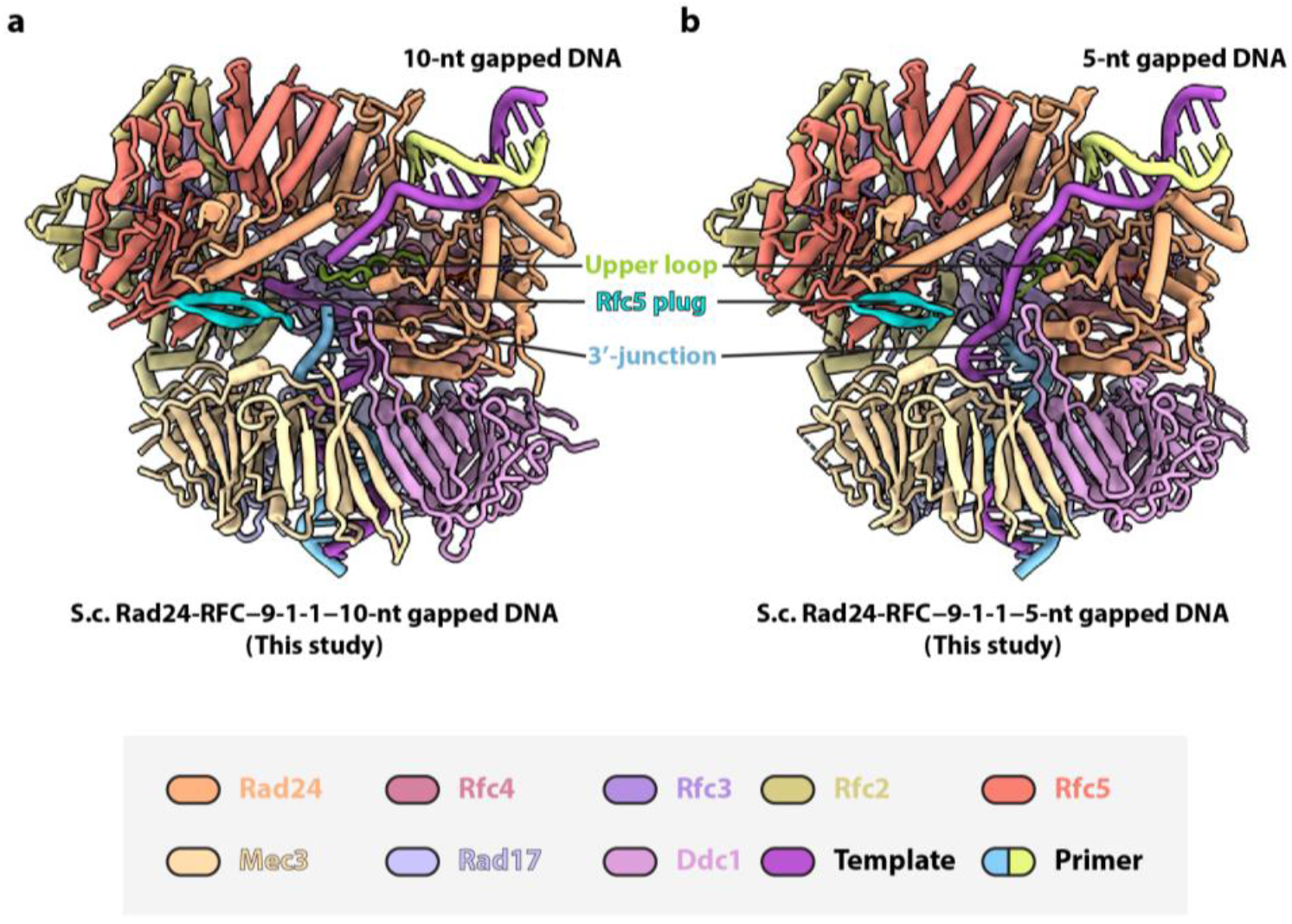
Comparison of 10-nt and 5-nt gapped DNA bound Rad24-RFC–9-1-1 structures both in the 9-1-1 clamp fully closed state. **a-b)** Structures of the 10-nt (a) and 5-nt (b) gapped DNA bound complexes. The key features including the Rad24 upper loop (green), the Rfc5 plug (cyan), and the 3′-DNA junction (blue) inside the loader chamber are labelled. The color scheme is shown at the bottom.

## LEGENDS FOR VIDEOS

**Video 1. Comparison of five loading intermediates of the 10-nt gapped DNA bound and one loading intermediate of the 5-nt gapped DNA bound Rad24-RFC–9-1-1 complexes.** The six atomic models are first rotated 360° around the Y axis, then rotated another 360° around the X axis. These models are aligned but shown separately.

**Video 2. An animation showing how the Rad24 upper loop blocks upward advance of DNA 3′-junction to enforce a minimum gap size.** First, the Rad24-RFC–9-1-1–10-nt gapped DNA and the RFC–PCNA–DNA structures are shown side by side rocking around the Y-axis. Then, the RFC complex is changed to transparent grey and superimposed onto the Rad24-RFC complex, followed by zooming into the center region of the structures. Next, the loaders disappear leaving only the 3′-end part of the primer strand and the loop harboring the separation pin residue (lime) for a better comparison. The superimposed models are rocked along the Y and X axes to better show the blocking of the DNA 3′-junction by the Rad24 upper loop (green).

**Video 3. Morphing of 3′-DNA entering the 9-1-1 clamp chamber.** Based on the five intermediate steps of the 10-nt gapped DNA bound complex captured in this study, the morph starts with the 5′-junction part of the gapped DNA already bound on the Rad24-RFC shoulder. First, the complex in step 5 is rotated 360° around the Y axis. Second, Mec3 is rotated 44° to the left to open a gap of 42 Å in the 9-1-1 clamp (also see Fig. 3). This gap is wide enough to allow passage of a dsDNA that is 20 Å wide. Third, the 3′-junction part of the gapped DNA gradually enters the 9-1-1 clamp chamber (as observed in steps 2 and 3). Fourth, when the DNA finishes entering, Mec3 rotates 44° to the right to close the 9-1-1 gate (as observed in steps 4 and 5). Same color scheme is applied here as in the main text figures.

